# HyperBind2: Multi-Shot Learning Enables Progressive Improvement in Computational Antibody Discovery

**DOI:** 10.1101/2025.11.06.687005

**Authors:** Daniel Dell’uomo, Andrew Satz, Brett Averso

## Abstract

Antibody discovery remains constrained by resource-intensive experimental screening approaches that offer limited control over critical properties. Here we present HyperBind2, a machine learning platform that progressively improves antibody-antigen interaction predictions through experimental feedback cycles. Unlike static or zero-shot computational approaches, HyperBind2 employs multi-shot learning, which adapts to target-specific patterns using minimal experimental data (10-20 validated binders/non-binders). The platform requires only the target’s primary sequences as input (no experimental structures are required), embedding antibody and antigen sequences into a shared representation space where binding affinity is modeled as a learned geometric relationship.

HyperBind2 was validated at multiple independent academic and commercial labs. One such experiment validated HyperBind2 on a challenging multi-pass membrane receptor target through three iterative design-test cycles. Starting from an initial screening of 100 million candidates completed within 48 hours, model accuracy improved from 65% to 85% across three rounds of lab-to-AI feedback. By round 3, HyperBind2 achieved a 21% experimental success rate, with 20 of 96 tested candidates demonstrating KD ≤ 100 nM, including 3 with sub-10 nM affinities. HyperBind2 spans multiple therapeutic formats including scFvs, VHHs, and full-length IgGs, with preliminary research extending to CAR-T, BiTE, and bispecific formats.

HyperBind2 establishes an efficient digital-experimental workflow that reduces laboratory resources and screening time while maintaining high hit rates. By combining massive computational pre-screening with targeted experimental validation and continuous model refinement, HyperBind2 significantly reduces experimental burden while accelerating the identification of therapeutic quality antibody candidates. HyperBind2 is available via open-source for academic research or a commercial platform (abtique.com), which provides lab-ready antibody sequences with no coding required.

**Disclaimer:** Due to the sensitive nature of intellectual property and confidentiality agreements, target identities have been anonymized where research is ongoing or proprietary. While we provide detailed descriptions of experimental methods, all details cannot be disclosed. The results have been independently validated by third parties, enhancing their reliability.

## 1. Introduction

### 1.1 The Challenge of Antibody Discovery

Antibody discovery remains largely reliant on animal immunization or high-throughput *in-vitro* screening, both of which are time-consuming, expensive, and offer limited control over specificity, diversity, and developability [1–12]. These methods frequently yield candidates with sub-optimal affinity, poor stability, or high immunogenicity, particularly when targeting novel or conformationally complex antigens [13–21].

While antibodies have become a cornerstone of modern therapeutics due to their ability to specifically recognize and bind to target molecules, significant challenges persist in their development. As of December 2024, over 200 antibody therapies have been approved and marketed [22], yet developing potent antibodies against challenging or novel targets remains a considerable hurdle [23–27].

These challenges have prompted interest in computational approaches, though current methods remain limited to comparatively low-throughput antibody modeling [28–33]. Even with these computational tools, traditional screening methods still face challenges where the immunogenicity of certain epitopes that are highly susceptible to immune recognition may complicate screening efforts, resulting in panels of antibodies with sub-optimal or immunodominant species ineffective for the desired mechanism of action [34–35].

This difficulty is compounded by the fact that antibody sequences are shaped by stochastic V(D)J recombination and somatic hypermutation (SHM) [36–38], resulting in highly diverse repertoires that are not directly predictable from sequence alone [39, 40]. Furthermore, antibody discovery pipelines rarely preserve information about true non-binders, making it difficult to train predictive models that can distinguish between positive and negative examples [41–45]. From a statistical perspective, this absence of validated negatives fundamentally limits the ability to estimate decision boundaries, calibrate confidence scores, and distinguish genuine biological signal from statistical noise, forcing models to learn from an incomplete representation of the underlying data distribution. Recent computational approaches have achieved impressive capabilities in antibody design, with some platforms reporting 15-20% success rates in initial validation studies. However, most of these reported results generated antibody designs to known targets with known antibodies with little evidence of antibody designs to unique, novel, or challenging targets. Given the lack of empirical evidence around single-shot design, the critical obstacle for difficult or novel targets lies not just in first-round results but in adapting to experimental feedback for progressive improvement.

While large generative models could theoretically incorporate experimental data through fine-tuning, practical implementation faces significant barriers. With billions of parameters but only tens to hundreds of experimental observations from any given campaign, determining appropriate training protocols becomes challenging—insufficient adaptation yields no improvement, while aggressive fine-tuning risks disrupting the model’s foundational capabilities. The computational resources required (extensive GPU time and specialized engineering) make project-specific adaptation economically feasible for only a handful of discovery teams. Additionally, therapeutic development requires strict data segregation between projects, creating a tension between the need for project-specific learning and the architectural realities of shared model infrastructures.

These constraints highlight why purpose-built multi-shot learning architectures offer practical advantages for iterative antibody optimization. Direct learning approaches can efficiently extract signal from limited experimental observations-transforming 96 tested sequences into thousands of training contrasts through synthetic amplification. This enables rapid model adaptation while maintaining complete data isolation between projects. Most importantly, multi-shot optimization naturally accommodates the high variability in success rates across different protein targets. Where a challenging membrane protein might yield only 5% initial success, the ability to learn from those failures and improve to 20% in subsequent rounds provides tangible value. Conversely, for well-studied targets (e.g. SARS-CoV-2, EGFR, PD-1) starting at 30% success, iterative refinement can push toward 50% or higher. This adaptive capability-learning what works for each specific target rather than applying static predictions-represents the key advantage of multi-shot learning over single-shot generation, regardless of the initial prediction quality. Multi-shot optimization aligns with the realities of drug discovery: limited experimental budgets, strict IP requirements, diverse target difficulties, and the need for continuous improvement rather than one-time predictions.

### 1.2 Computational Antibody Design: Promise and Gaps

Recent advances in computational protein design have shown promise, but significant challenges remain for antibody engineering. Diffusion-based protein-design methods (e.g. RFdiffusion [46]) have demonstrated success for globular proteins but struggle with antibody-specific structural idiosyncrasies and the requirement for precise epitope targeting [47, 48]. Moreover, these structure-based approaches typically rely on static crystallographic or cryo-EM snapshots that fail to capture the inherent conformational dynamics of antibodies, which undergo significant structural rearrangements during binding and can sample multiple functional states[49]. Protein language models (PLMs) encode biologically meaningful features (such as somatic hypermutation (SHM) burden, isotype class, and paratope features) but they are typically trained on static databases and do not capture the dynamic selection forces active *in vivo* [50–53]. Furthermore, antibody specificity is context-dependent and often structurally ambiguous, making it difficult to infer paratope-epitope relationships from sequence alone using general-purpose models [54–58].

Computational protein design (’de novo’ design) has been a topic of scientific inquiry for decades but has only recently begun to demonstrate practical feasibility for large, complex molecules such as antibodies or novel protein scaffolds [59–62]. The challenge of effectively designing antibodies for challenging drug targets is that there is very limited or no training data available for most novel targets [63–65].

The field has witnessed a proliferation of computational and machine learning methods aimed at predicting antibody-antigen interactions through both sequence-based and structural approaches [28]. These include tools for paratope prediction [66–68], epitope mapping [69], and comprehensive paratope-epitope interaction modeling [70–73]. While this expanding toolkit shows promise (with paratope prediction generally outperforming epitope prediction in accuracy [74]), a fundamental question remains unresolved: whether antibody-antigen interactions can be reliably predicted from first principles, and if so, what theoretical framework would underpin such predictions [30, 75, 76].

Several key limitations persist in current computational approaches:

– **Structural dependency**: Most methods require high-quality structural data, limiting their applicability to challenging targets like GPCRs, ion channels, and intracellular proteins [77–81].
– **Limited training data**: Sparse experimental binding data, particularly for true negative examples, constrains model training and generalization [82–85].
– **Poor transfer learning**: Models trained on one target often fail to generalize to new targets without extensive retraining [86–89].

#### 1.2.1 The Zero-Shot to Multi-Shot Evolution

Breakthroughs in zero-shot antibody design have demonstrated impressive capabilities, but reveal telling patterns in target selection. RFdiffusion validated designs against influenza hemagglutinin, IL7Rα, insulin receptor, and PD-L1 all established therapeutic targets with dozens of known antibodies and extensive structural characterization. Their antibody design work focused on influenza HA, RSV site III, SARS-CoV-2 RBD, and C. difficile toxin B, each having hundreds of published antibody sequences and structures. Absci concentrated on HER2 (the target of Herceptin/trastuzumab), VEGF-A, and SARS-CoV-2 Omicron variant-proteins with thousands of documented binders and billions in pharmaceutical development. Even ESM-1v’s mutation prediction validated on proteins like GFP, UBC9, and protein G, which have been studied for decades with comprehensive mutational landscapes mapped. This pattern suggests zero-shot methods appear to be limited to adapt to well characterized targets where prior antibody data exists, minimizing breakthroughs for novel, challenging, or ‘undruggable’ proteins.

#### 1.2.2 The Reality Gap: Zero-Shot on Novel Targets

Real antibody development faces cascading failure modes that zero-shot methods fundamentally cannot predict. From initial libraries, variants form insoluble inclusion bodies at 37°C, aggregate below therapeutic stability requirements of 65°C, fail biophysical screens for viscosity and polyreactivity, or show no functional activity. These failures stem from complex, interrelated factors: glycosylation that can either enable or destroy epitope recognition with patterns varying drastically between proteins; conformational dynamics invisible to structural methods; and conflicts between assays where strong SPR binding becomes complete failure in cell-based assays. For heavily glycosylated proteins, the conformational entropy loss from constraining glycans during antibody binding creates thermodynamic barriers that make high-affinity recognition nearly impossible, while glycan shielding blocks access to critical epitopes entirely.

Most zero-shot methods on well-characterized targets with documented antibodies and comprehensive structural data creates an illusion of capability. These models essentially interpolate within dense clouds of existing knowledge, having seen hundreds of similar binding modes and epitope patterns during pre-training. This fundamentally differs from approaching truly novel targets where glycosylation patterns remain unknown, conformational states unmeasured, and no similar examples exist in any database. For genuinely challenging proteins heavily glycosylated checkpoints, dynamic GPCRs, or intrinsically disordered proteins zero-shot methods cannot predict whether epitopes will be glycan-blocked, whether expression system differences will alter conformations, or how binding requirements interact with stability, viscosity, and polyreactivity constraints. This explains why pharmaceutical companies still struggle with “undruggable” targets despite computational advances: these proteins exist in sequence-function space where no training data exists, making zero-shot extrapolation impossible.

### 1.3 Our Approach: Hyperbind2

To address these challenges, we introduce **Hyperbind2**, a platform based on contrastive learning [90–93] for de-novo antibody screening and engineering, using a multi-shot learning architecture with four key components:

1. **Large foundational embedding models** Proprietary antibody structure and sequence encoders, and protein structure and sequence encoders (e.g., ProtBertBFD), providing consistent protein representations that remain frozen during project-specific learning
2. **Synthetic amplification of experimental outcomes** that transforms limited observations (e.g., 96 tested sequences) into comprehensive training signals (∼9,000 contrastive pairs) through biologically-informed pairing strategies, extracting maximum learning from both successful and failed designs
3. **Direct learning architecture** with lightweight adaptation layers trained exclusively on user’s experimental data, maintaining complete data segregation between projects while enabling efficient learning from 10-100 experimental observations
4. **Iterative feedback mechanism** enabling 24-48 hour model updates between experimental rounds, creating compound learning effects where each round’s outcomes inform subsequent predictions

While the foundational encoder incorporates both sequence and structure-derived features during pre-training, inference requires only sequence input, eliminating dependencies on crystal structures or AlphaFold predictions [94]. Hyperbind2 jointly embeds antibody and antigen sequences into a learned representation space, modeling binding affinity as a geometric relationship rather than requiring explicit structural modeling or docking.

Hyperbind2 is not a generative model; it does not generate antibody structures or produce antibody-antigen complexes [95]. Instead, it ranks and prioritizes existing sequence candidates from large libraries (10^8-10^9 sequences) based on predicted binding likelihood. The platform uses protein language modeling to encode biophysical properties-CDR geometry, flexibility patterns, hydrophobicity distributions-as latent features within the embedding space [39, 96, 97].

A key innovation is the platform’s ability to learn from complete experimental outcomes, including the 80-90% of designs that typically fail validation. Through synthetic amplification, each experimental round generates thousands of training signals from both positive and negative results, enabling robust model adaptation from campaigns as small as 96 sequences per round. This multi-shot learning approach has demonstrated progressive improvement in our chemokine receptor campaign: Round 1 yielded 5-10 weak binders (100-1000nM), Round 2 achieved 3 sub-100nM binders, and Round 3 produced 20 sub-100nM binders including a 0.84 nM lead-representing progression from <10% to 21% success rate within three iterative cycles.

The platform currently screens 10-100 million antibody designs in under 48 hours and supports diverse therapeutic formats including VHHs, scFvs, and full-length IgGs, with validation extending to complex formats including bispecifics.

#### Implementation Strategy

We provide two implementations serving different communities:

1. **Open-Source Version**: Built on ESM3 encoder [98] with a contrastive learning “head” architecture, designed for academic research and methodological transparency. This version democratizes access to multi-shot learning capabilities for antibody design.
2. **Commercial Platform**: Utilizes proprietary encoders and advanced features developed for our commercial partnerships, providing enhanced performance, automated workflows, data engineering pipelines, and complete data segregation for therapeutic development programs. Visit abtique.com for access.

By enabling efficient learning from limited experimental observations through synthetic amplification and multi-shot optimization, Hyperbind2 transforms antibody discovery from static prediction to progressive improvement, potentially accelerating therapeutic development while reducing experimental burden.

## 2. Methods

### 2.1 Overview of Hyperbind2 Workflow

Hyperbind2 employs a systematic computational-experimental pipeline designed to identify high-affinity antibodies from vast sequence spaces with minimal structural input requirements. The platform operates through five sequential stages that combine high-throughput *in silico* screening with targeted experimental validation:

**Figure 1.**
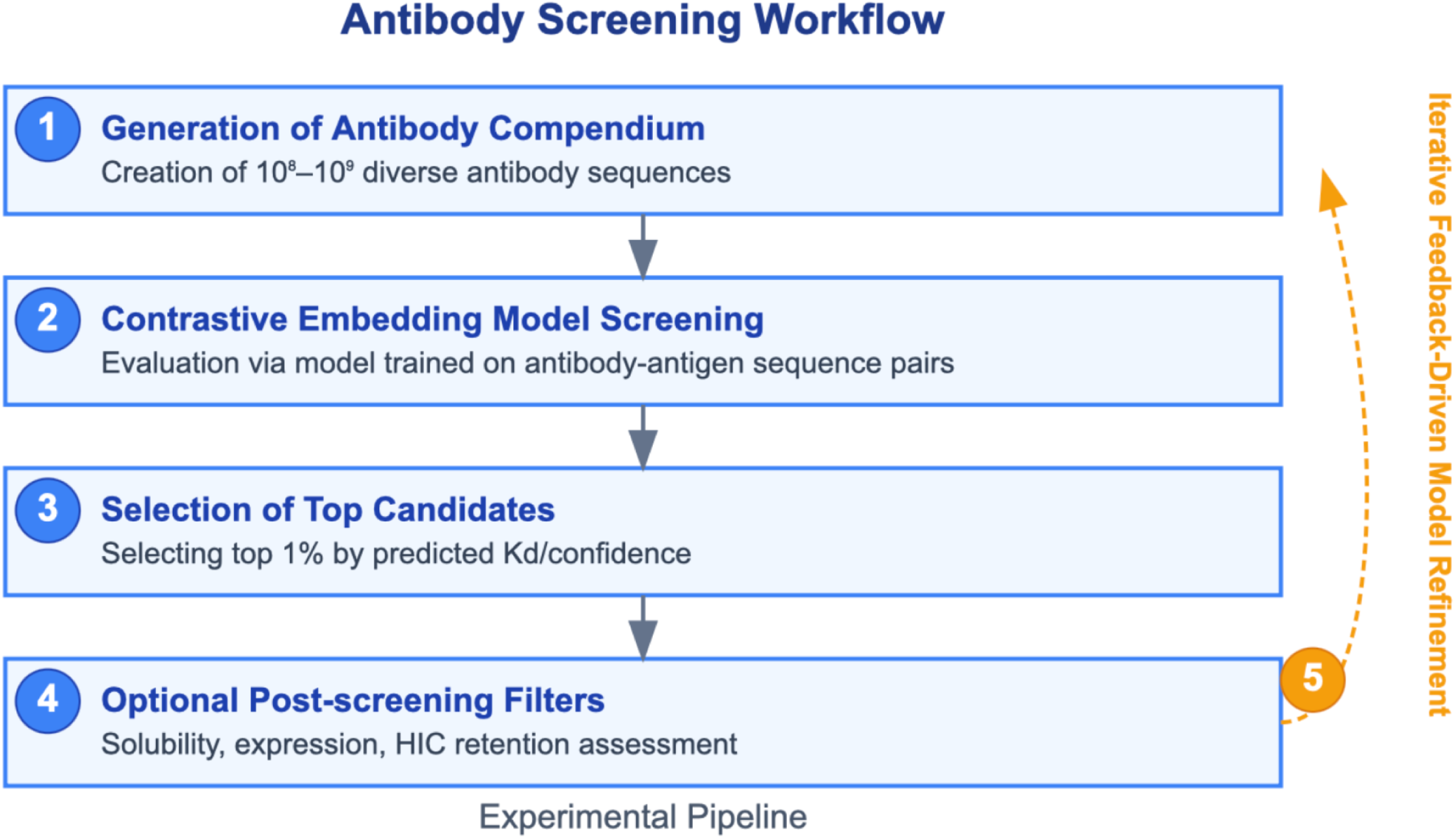
Schematic overview of the computational antibody screening workflow. The pipeline consists of five sequential stages: (1) generation of a diverse antibody sequence compendium (10⁸–10⁹ sequences); (2) in silico screening via a contrastive embedding model trained on antibody-antigen sequence pairs; (3) selection of top-ranked candidates (top 1% by predicted Kd/confidence); (4) application of optional post-screening filters to assess biophysical properties (solubility, expression, HIC retention); and (5) experimental validation with iterative feedback to the computational model. This computational-experimental framework enables efficient screening of vast antibody repertoires while continuously refining predictive accuracy through experimental validation.

This computational-experimental framework enables efficient screening of vast antibody repertoires while continuously refining predictive accuracy through experimental validation. The workflow integrates *in silico* hit prediction with *in vitro* validation for antibody binding and characterization, enabling the screening of over a million antibody sequences in a single hour.

**Figure 2:**
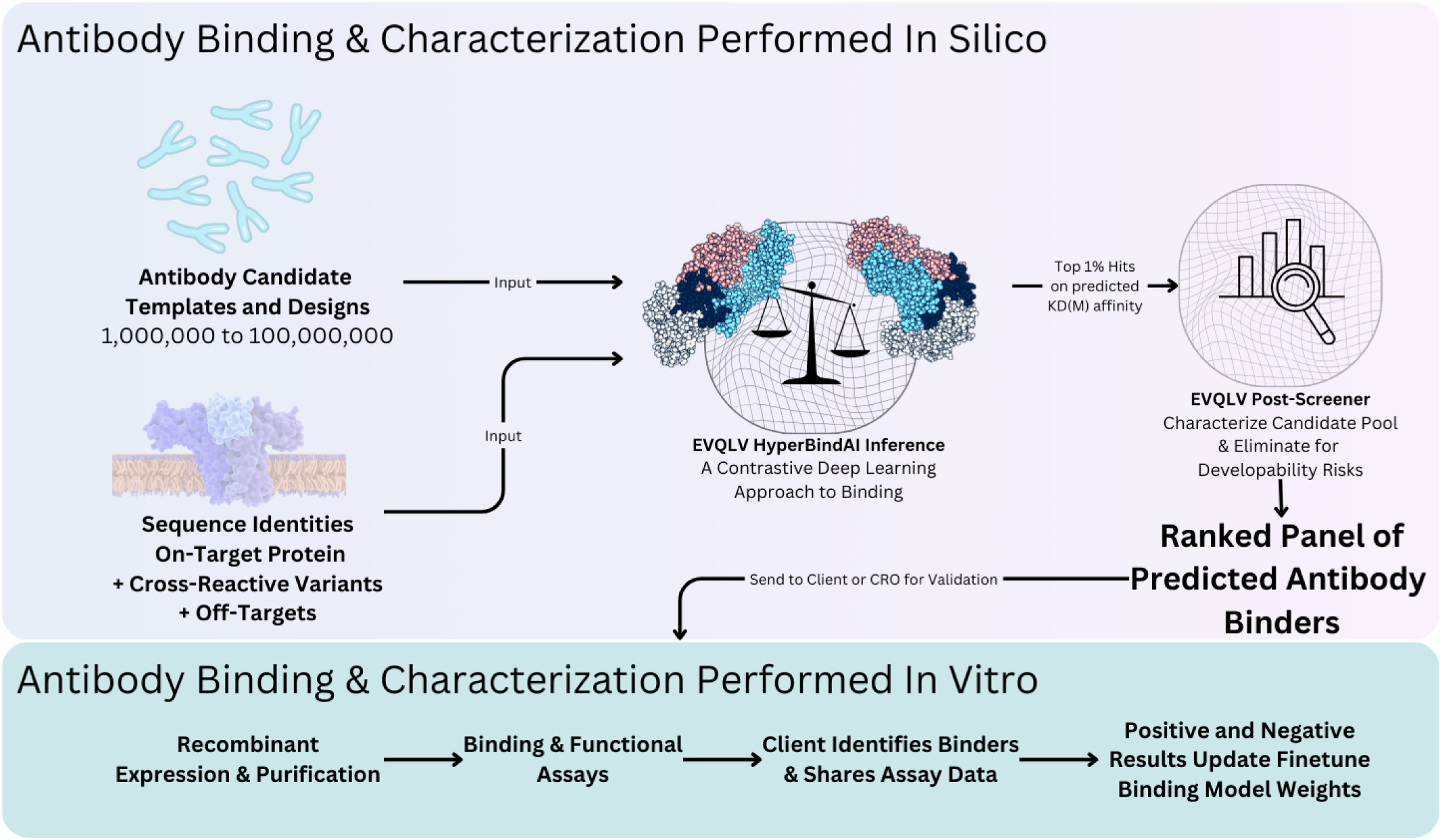
Antibody Binding & Characterization Performed In Silico. From in silico prediction to in vitro validation: a streamlined workflow for antibody discovery involves multi-shot learning cycles of digital inference combined with wet lab efforts for binding and characterization. Each iteration incorporates both positive and negative outcomes to refine and update model learning, driving continuous improvement in antibody design and optimization.

### 2.2 Antibody Sequence Generation Platform

The Hyperbind2 platform employs digital methodologies to generate novel antibody and nanobody fragment variable (Fv) sequences from scratch. By harnessing advanced machine learning techniques, EVQLV creates unique Fv sequences that are tailored for both therapeutic and research needs. Several machine learning-based approaches are leveraged to create digital compendiums of antibody sequences which include:

– **VDJ recombination simulation**: Modeling the natural process of V(D)J recombination [38] to generate biologically plausible antibody sequences that recapitulate natural diversity patterns
– **Masked-Language In-Filling**: Using protein language models [99] to fill in masked regions of antibody sequences while maintaining structural and functional constraints
– **Autoencoding**: Learning compressed representations of antibody sequences for efficient generation of novel variants [100]
– **Fv Structure-Based Diffusion**: Generating sequences based on structural constraints and diffusion processes that preserve key geometric features [101]
– **Inverse Folding**: Designing amino acid sequences that fold into target antibody structures by learning sequence-structure relationships from known proteins [102]

**Figure 3:**
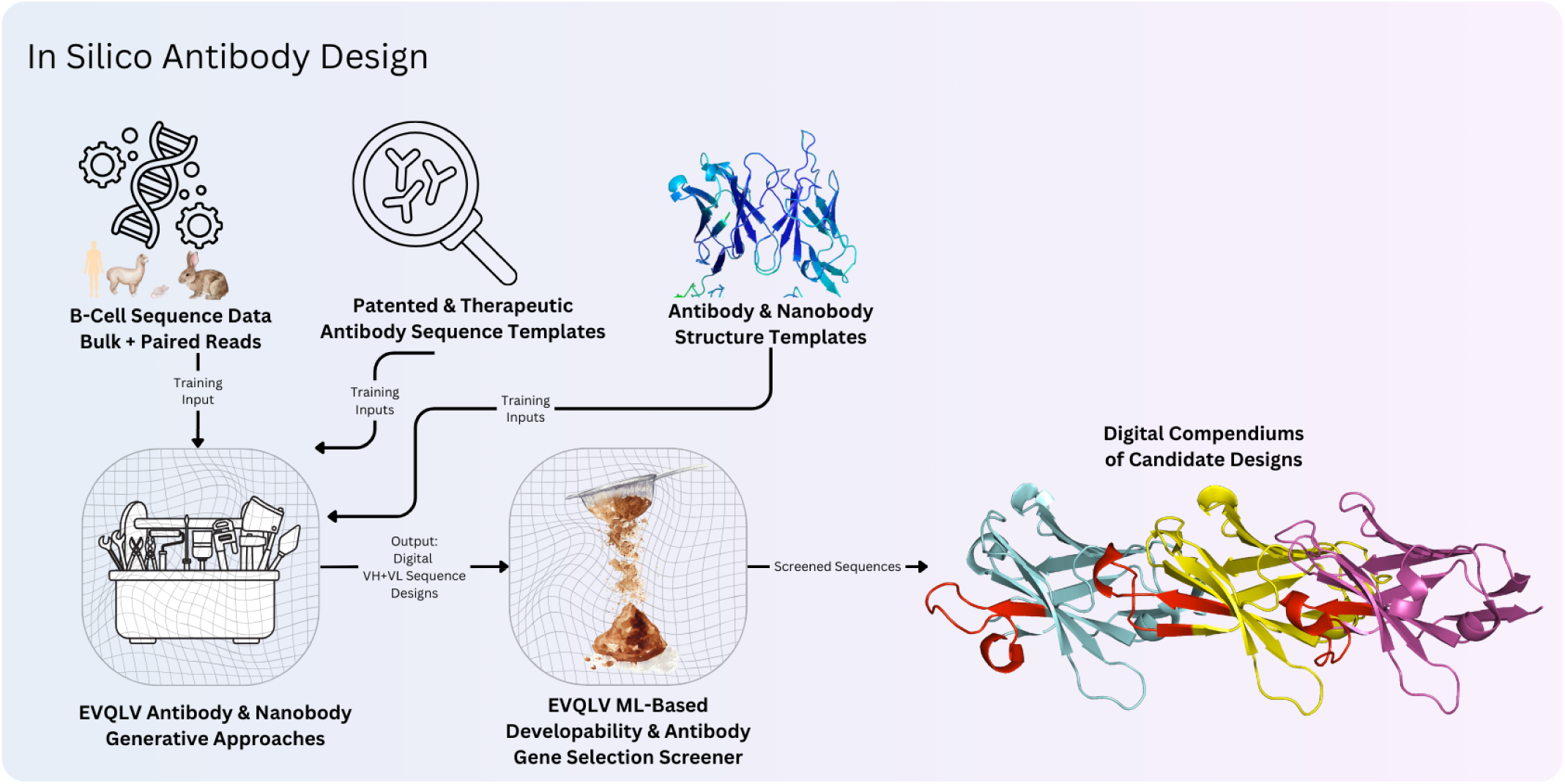
In Silico Antibody Design Methods. Each model has unique training input requirements, utilizing both public and licensed data sources. Machine learning and deep learning approaches are applied to generate a comprehensive antibody compendium. The model outputs are stored in a database and pre-screened for developability and desirable Ig gene characteristics [8]. Together, these methods enable a diverse and robust computational framework for designing antibody and single-domain antibody Fvs.

Additional screening is applied subsequently to select for developable candidates before storing into a digital compendium. Critical factors considered at this stage include hydrophobicity profile and solubility predictions, CDR length distributions and structural feasibility, heavy-light chain pairing compatibility, immunoglobulin germline gene compatibility factors, and expression likelihood and aggregation propensity analysis [8].

Several digital compendiums have been constructed using a variety of approaches, with a complete population of over 3 billion pre-screened paired sequence designs to date. Each model has unique training input requirements, utilizing both public and licensed data sources, with machine learning and deep learning approaches applied to generate a comprehensive antibody compendium.

### 2.3 Hyperbind2 Architecture: Foundation Model to Direct Learning

#### 2.3.1 Foundation Model Training

##### Dataset Assembly and Curation

The foundation of Hyperbind2’s predictive capability begins with pre-training on an extensive dataset representing over five years of systematic curation. Unlike publicly available databases containing primarily successful therapeutic antibodies, the dataset uniquely captures the full spectrum of binding outcomes-from picomolar binders to confirmed non-binders-providing the negative examples essential for training discriminative models.

The foundation dataset aggregates information from multiple sources:

– **Patent Literature Mining**: Systematic extraction from global patent databases (2000-2024), capturing marketed therapeutics, optimization trajectories, and negative controls from experimental sections
– **Scientific Literature Curation**: Analysis of tens of thousands of publications, particularly supplementary materials where negative results are often reported but rarely indexed
– **Proprietary Data Partnerships**: Complete experimental campaigns, including the variants that typically remain unpublished

This diversity spans multiple antibody formats (IgG, scFv, Fab, VHH, bispecifics), assay types (SPR, BLI, ELISA, flow cytometry), and including both successes and failures, provides critical advantages over models trained solely on positive examples from the Protein Data Bank [103] or therapeutic databases [104].

##### Addressing Foundation Dataset Limitations Through Synthetic Generation

Despite extensive curation, even our foundation dataset faces limitations requiring synthetic augmentation:

– **Limited true negative examples**: Discovery pipelines rarely preserve non-binders, creating imbalanced training data
– **Mutational sensitivity**: Single mutations can eliminate binding, requiring dense sampling around known binders
– **Dataset bias**: Public databases skew toward human therapeutic scaffolds
– **Target coverage gaps**: Most targets have <100 validated binders; novel targets may have none

To address these limitations, we augment the foundation dataset with synthetic sequences generated from experimentally validated binders across multiple targets including HIV, coronavirus, and influenza antibodies, plus entries from the Immune Epitope Database (IEDB) [105].

Synthetic generation follows two strategies:

1. **Local edits** (subtle variants maintaining binding likelihood):

– 1-3 mutations from parent sequences [106]
– CDR RMSD < 1.5 Å when structures available [107]
– Embedding similarity > 0.9 to parent
2. **Global rewrites** (negative examples with structural plausibility):

– Preserved framework architecture
– Embedding similarity < 0.8 to ensure diversity
– Retained as “non-functional expressibles” after biophysical screening [108]

This synthetic augmentation provides the foundation model with comprehensive positive-negative training examples before any user-specific adaptation occurs.

##### Foundation Encoder Architecture

Using this comprehensive dataset, we pre-train proprietary encoders that generate consistent, high-dimensional embeddings capturing both sequence and structural features:

– **Antibody Encoder**: Processes VH/VL sequences, learning CDR geometry, framework compatibility, and developability features
– **Antigen Encoder**: Embeds target sequences, capturing epitope accessibility and binding site characteristics
– **Training Objective**: Contrastive learning on millions of known binder/non-binder pairs from the curated dataset
– **Output**: 1280-dimensional embeddings per sequence that remain frozen during user-specific adaptation

These foundation encoders provide the stable representational backbone that enables efficient learning from limited user data.

#### 2.3.2 Direct Learning Architecture for User-Specific Adaptation

While the foundation model provides general protein representations, therapeutic discovery requires adaptation to specific experimental systems. Our direct learning architecture achieves this through a lightweight head model that trains exclusively on user data:

##### Architectural Separation

– **Foundation Layer (Frozen)**: Pre-trained encoders providing consistent embeddings
– **Direct Learning Head (Trainable)**: Project-specific layers learning from user’s experimental outcomes

##### Direct Learning Head Structure

– Input: Concatenated antibody-antigen embeddings (2560 dimensions)
– Hidden layers: [2560→1024→512→256] with ReLU activation
– Output: Binding affinity prediction (scalar) + confidence score
– Parameters: ∼2.7M trainable weights (vs billions in foundation model)

##### Learning Protocol

– Round 1: Initialize with generic weights
– Post-Round 1: Update using experimental results (e.g., 10 binders + 86 non-binders)
– Round 2 to Round N: Incremental updates with accumulated data
– Complete data segregation: Each project maintains isolated weights

This separation ensures the foundation model’s general knowledge remains intact while enabling rapid, project-specific learning that adapts to the project-specific target, expression system, and assay conditions.

#### 2.3.3 Synthetic Amplification: Transforming Limited Data into Comprehensive Training

Despite the foundation model’s extensive pre-training, user-specific adaptation requires learning from just 10-100 experimental observations. Our synthetic amplification pipeline transforms these limited observations into thousands of informative training pairs.

##### Data Acquisition Through Modified Protocols

Obtaining sequence information from experimental failures requires significant deviation from conventional laboratory protocols. Standard procedures discard wash fractions, expression failures, and below-threshold measurements without sequence determination. We have developed modified protocols that systematically capture and sequence these conventionally discarded materials-modifications detailed in a pending patent application. The key point is that accessing negative experimental data requires intentional changes to standard workflows.

##### Four Methods of Synthetic Amplification

###### 1. Homologous Pairing with Divergent Outcomes

We identify sequence pairs with high embedding similarity (cosine scores 0.90-0.99) but opposite experimental outcomes. For example, two variants differing by only 2-3 CDR residues where one exceeds a binding threshold (KD <100nM) and another fails to meet the threshold or is beyond the limits of detection, providing gradient information about critical binding determinants. Each successful binder is systematically paired with 10-20 similar failures, creating focused contrasts that teach the model precise decision boundaries rather than broad sequence patterns.

###### 2. Hard Negative Mining Near Decision Boundaries

Sequences with borderline experimental outcomes (∼KD 500nM) undergo targeted perturbation to generate challenging negative examples. Using BLOSUM62 substitution matrices, we introduce unfavorable mutations (scores <-2) at CDR positions, creating variants predicted to fail while maintaining structural plausibility. Additionally, gradient-based analysis in the embedding space identifies the steepest descent directions toward non-binding regions. This generates 20-30 hard negatives per borderline sequence, forcing the model to learn subtle discriminative features rather than obvious sequence differences.

###### 3. Systematic Perturbation Protocols

All experimental sequences undergo controlled modifications to explore the functional landscape:

– **CDR swapping**: Exchange CDR regions between failed variants to identify incompatible combinations
– **Domain shuffling**: Recombine framework and CDR elements from different experimental outcomes
– **Motif deletion**: Remove suspected binding motifs to verify their necessity
– **Progressive mutation**: Introduce 1-5 mutations along paths from binders to non-binders

Each experimental sequence generates 50-100 synthetic variants through these perturbations, with labels interpolated based on the severity and location of modifications.

###### 4. Adaptive Weighting by Information Content

Training pairs receive differential weights based on rarity and informativeness:

– Expression failures (appearing in <5% of sequences): 5× weight multiplication
– Rare binding modes: 3× weight
– Common failure patterns: 0.5× weight
– Near-threshold variants: 2× weight

This ensures the model learns from the most informative examples rather than being dominated by abundant but uninformative failures.

#### 2.3.4 Contrastive Learning Framework

The contrastive learning objective operates at both foundation and direct learning stages:

##### Foundation Model Contrastive Training

– Millions of known binder/non-binder pairs from curated dataset
– Learn general antibody-antigen interaction patterns
– Create embedding space where binding correlates with proximity

##### Direct Learning Contrastive Adaptation

– User’s specific experimental outcomes
– Refine embedding relationships for specific target
– Maintain general knowledge while learning target-specific patterns

The model is trained such that experimentally validated binder pairs experience attractive forces within the embedding space, while non-binders are systematically repelled. This creates a high-dimensional landscape enabling rapid ranking of candidate antibodies against target antigens [94].

#### 2.3.5 Integration with Multi-Shot Learning

The synthetic amplification pipeline operates iteratively across experimental rounds through selective pairing:

– **Round 1**: Initial 96 sequences → ∼1,800 informative training pairs
– **Round 2**: Accumulated 192 sequences → ∼3,600 cumulative training pairs
– **Round 3**: Accumulated 288 sequences → ∼5,400-9,000 cumulative training pairs

While synthetic amplification could theoretically generate O(n²) pairs (96 sequences could yield ∼9,000 total pairs), we employ selective pairing focusing on the most informative contrasts-sequences with high similarity but opposite outcomes, near-threshold variants, and rare failure modes. This selective approach prioritizes quality over quantity, ensuring each training pair provides maximum gradient information for model improvement.

Each round’s amplified data incorporates patterns discovered from previous rounds, creating compound learning effects. The model doesn’t just memorize which sequences worked-it learns what patterns to avoid, enabling the progressive improvement from <1% to 23.9% success rate demonstrated in one case study highlighted below.

This synthetic amplification methodology, combined with the contrastive learning architecture, transforms the fundamental economics of antibody discovery: where traditional approaches require screening millions of variants to find viable candidates, our approach achieves comparable or superior results with less than 300 designed sequences by extracting maximum learning from every experimental outcome.

### 2.4 Structure-Agnostic Design for High-Throughput Screening

A key practical advantage of Hyperbind2 lies in its structure-agnostic architecture-the model can directly encode structural information when provided but operates effectively on sequence alone, enabling screening of millions of antibody candidates without computational bottlenecks.

The foundation encoders demonstrate remarkable flexibility in input handling. When provided with sequences alone, the encoder generates structure-aware embeddings based on patterns learned during extensive pre-training, capturing implicit structural features without explicit 3D coordinates. Alternatively, when structural data is available-whether from AlphaFold2/3, crystal structures, or homology models-the encoder directly incorporates these 3D coordinates into the embedding representation. In practice, we typically adopt a hybrid approach: providing the antigen structure, which needs to be encoded only once for the entire screening campaign, while evaluating millions of antibody sequences without requiring their individual structures. This asymmetric strategy leverages structural information where it’s most valuable and computationally feasible while maintaining throughput for massive antibody libraries.

The practical impact of this design becomes clear when examining computational requirements. In a standard screening campaign, we encode the antigen structure from AlphaFold or PDB in approximately 10 seconds-a one-time computational investment. We then screen 10^8 antibody sequence variants against this target, completing the entire evaluation within 48 hours without generating a single antibody structure. If structural prediction were required for each antibody candidate, the same campaign would require months of computation, as AlphaFold predictions take 2-5 minutes per sequence. The encoder achieves this efficiency by learning during foundation training to map antibody sequences directly to structure-aware representations that capture CDR geometries, framework orientations, and binding interface properties, all without explicit structure prediction.

This structure-agnostic operation proves particularly valuable for challenging therapeutic targets. For membrane proteins like GPCRs that resist crystallization, we can screen antibody libraries without waiting for elusive structural data. When pursuing novel or emerging targets, optimization campaigns can begin immediately using sequence information while structural characterization proceeds in parallel. The approach also handles conformationally flexible proteins effectively, as the learned embeddings capture sequence-structure relationships that aren’t confined to single static conformations.

Most critically, this design enables the rapid iterative cycles that define our multi-shot learning approach. Processing millions of candidates per round, incorporating experimental feedback, and generating updated predictions within 24-48 hours forms the backbone of our progressive optimization strategy. If AlphaFold structure prediction were required for each antibody variant, this rapid iteration would be impossible. The ability to operate on sequence alone while maintaining structure-aware representations through learned embeddings enabled the dramatic improvement from less than 10% to 21% success rate demonstrated in our chemokine receptor campaign, validating that computational efficiency and biological accuracy can be achieved simultaneously.

### 2.6 Multi-Shot Learning Protocol

#### Round 1 - Initial Exploration

– Screen 10^6 - 10^8 candidates using base model (proprietary antibody sequence library input)
– Ranking based on structure-aware embeddings learned during foundation training
– Select 96 diverse candidates (uncertainty sampling + clustering)
– Experimental validation via SPR/BLI
– Data collected: binding kinetics (KD values), expression levels, biophysics assay measures
– Typical outcome: 0 to 15% success rate

#### Inter-Round Learning

– Extract successful binders and non-binders from experimental data
– Generate contrastive training pairs through selective amplification: 96 experiments → ∼1,800 high-information pairs
– Fine-tune prediction head model (3 epochs, learning rate 1e-05)
– Re-rank entire candidate pool with updated, target-specific model

#### Round 2 - Focused Exploitation

– Prediction scores show ∼30% better separation between binders/non-binders
– Model has learned target-specific sequence patterns from Round 1 outcomes
– Select 96 candidates from refined high-confidence regions
– Experimental validation
– Cumulative learning from 192 total experiments
– Typical outcome: 15-25% success rate

#### Convergence Assessment

– Monitor accuracy improvement: Δaccuracy < 5% indicates convergence
– Typical convergence: 2-5 rounds (empirical range from multiple campaigns)
– Final model incorporates 200-500 experimental data points
– Training pairs after synthetic augmentation (96 experiments → ∼1,800 pairs per round, so 3,600-9,000+ pairs total)
– Target-specific model learns epitope-relevant structural features from experimental feedback

### 2.7 Commercial Implementation

Our proprietary commercial platform employs advanced custom encoders specifically designed for antibody-antigen interactions. The commercial version incorporates several enhancements that cannot be disclosed due to intellectual property considerations, including specialized antibody encoders trained on proprietary datasets, advanced attention mechanisms for improved paratope-epitope modeling, enhanced confidence scoring through ensemble methods, and optimized inference pipelines for large-scale screening applications using NextFlow [109]. The commercial platform, designed specifically for antibody scientists, is available at https://abtique.com/

## 3. Results

We demonstrate Hyperbind2’s transformative capabilities through a comprehensive real-world case study targeting a chemokine receptor—a notoriously challenging multi-pass membrane protein where traditional methods have historically failed. Starting from computational predictions with <10% success rate, we achieved 21% success rate with 20 sub-100nM binders and a best-in-class 0.84 nM affinity within just 6 months and 288 experimental tests. This improvement—accomplished through multi-shot learning that progressively refines predictions based on iterative experimental feedback—validates our approach of learning from complete experimental outcomes to achieve what traditional screening of millions of variants could not accomplish.

### 3.1 Model Performance and Synthetic Dataset Validation

The Hyperbind2 model was trained and evaluated using a combination of real-world and synthetic antibody datasets using a proprietary synthetic generation pipeline.

For development purposes, composite pairs based on antigen and binding-class information were created, generating both positive pairs (antibodies that bind to the same antigens) and negative pairs (antibodies targeting different antigens). This composite-pairing strategy significantly improved model performance compared with random pairing, increasing ROC-AUC from ∼0.53 to ∼0.76.

#### Training Efficiency and Convergence

The model demonstrated rapid convergence during training, reaching optimal performance within 2 epochs (∼36 minutes total training time). This rapid convergence demonstrates the effectiveness of the contrastive learning approach and suggests that the model efficiently captures the underlying binding relationships in the training data without requiring extensive computational resources.

#### Performance Metrics

A comprehensive evaluation revealed strong predictive capability across multiple metrics:

– **ROC-AUC**: 0.76 on held-out test sets, demonstrating robust discrimination between binders and non-binders
– **Precision-Recall curves**: Optimal F1-scores achieved through validation threshold optimization
– **Cross-validation**: Five-fold cross-validation demonstrated consistent performance across data splits

### 3.2 Computational Screening and Platform Capabilities

The platform demonstrates remarkable computational efficiency and scalability:

#### Screening Performance

– Screening over 1 million antibody sequences per hour during peak operation
– 10⁸ designs ranked in under 48 hours for large-scale screening projects
– Digital compendium of over 3 billion pre-screened paired sequence designs available for screening
– Top 1% candidates selected based on predicted Kd affinity and model confidence scores

#### Post-Screening Assessment

High-scoring candidates undergo further scrutiny with developability risk assessment, including direct prediction of critical biophysical properties such as SDS-PAGE profiles and monodispersity, hydrophobic interaction chromatography (HIC) retention prediction, expression levels and solubility predictions, and aggregation propensity and thermal stability metrics [110].

### 3.3 Case Study: De Novo Design of scFv Binders to Chemokine Receptor

We present a comprehensive case study demonstrating Hyperbind2’s capabilities in a challenging real-world application targeting a chemokine receptor [111] involved in immune cell signaling.

#### 3.3.1 Design Objectives and Target Specifications

The target product profile specified challenging requirements that traditional methods had failed to achieve:

##### Primary Objectives

– De novo design scFv antibodies targeting a specific chemokine class receptor involved in immune cell signaling
– Achieve high-affinity binding (KD ≤ 100 nM) with the aim of identifying multiple candidates exhibiting sub-10 nM affinities
– Ensure candidates exhibit high solubility, stability, and low aggregation propensity for therapeutic development suitability
– Demonstrate potential for modulation of receptor-mediated signaling pathways

###### Target Characteristics

The target receptor represents a challenging case for computational design due to its multi-pass membrane topology [112], limited high-quality structural information, and the requirement for functional modulation rather than simple binding inhibition.

#### 3.3.2 Design Approach and Compendium Generation

##### Foundation Dataset

The design process began with creation of a digital compendium using a proprietary autoencoder architecture, trained on a dataset of 2.1 million paired human VH+VL sequences to capture both sequence-level and structural features unique to antibodies.

##### Ground Truth Integration

This framework leveraged a ground truth dataset of 8 experimentally validated binders with high affinity (KD ≤ 10 nM) to various members of the chemokine receptor family, forming the foundation for target-specific optimization.

##### Compendium Scale and Diversity

A digital compendium comprising 6 × 10⁸ sequence candidates was generated, each systematically designed to expand upon the diversity of the ground truth binders while preserving structural relevance to the target receptor. Key metrics included:

– VH gene identity to nearest germline: 70-85% (IGHV1 and IGHV3 families used exclusively on heavy chains)
– VL gene identity to nearest germline: 85-95%, maintaining compatibility for downstream expression

#### 3.3.3 Multi-Shot Learning Progression and Quantitative Analysis

##### Empirical Learning Metrics Across Rounds

###### Round 1 Baseline

– Computational selection: 96 highest confidence predictions from 6 × 10⁸ candidates
– Experimental reality: 5-10 weak positive signals (KD 100-1000nM) out of 96 tested
– Majority showed KD >10³ nM despite high computational confidence scores
– 11/96 expression failures (inclusion bodies/aggregation)
– Key finding: Gap between computational confidence and experimental reality

###### Round 2 After Learning

– 100% expression success (vs 88% in Round 1)
– Several candidates achieving KD 10²-10³ nM
– 3 candidates <100nM (substantial improvement from Round 1)
– Key learnings applied: CDR-H3 length constrained to 10-15 amino acids, hydrophobicity (GRAVY) maintained between −0.2 to 0.3
– Updated model incorporated failure patterns from Round 1 to redefine “high confidence” predictions

###### Round 3 Convergence

– 20 candidates achieved KD ≤100 nM (21% success rate)
– 3 candidates demonstrated sub-10 nM affinities
– Best candidate: 0.84 nM binding affinity
– Marked reduction in non-specific binding and low-concentration artifacts
– Near-complete elimination of expression failures

##### Information Gain Through Synthetic Amplification

The method transforms limited experimental observations into comprehensive training signals through selective pairing and synthetic amplification:

– Round 1: 96 sequences tested → ∼1,800 high-information training pairs through selective contrast generation
– Round 2: 192 cumulative sequences → ∼3,600 cumulative training pairs
– Round 3: 288 cumulative sequences → ∼9,000 cumulative training pairs
– Training efficiency achieved through selective pairing of sequences with high similarity but divergent outcomes

###### Round 1 - Initial Screening

– Selection from the comprehensive compendium of 6 × 10⁸ paired VH/VL designs
– Hyperbind2 employed to predict binding affinity (KD ≤ 100 nM target)
– 96 scFv designs selected as highest computational confidence scores for experimental validation
– These top-scoring predictions served as a critical test of the foundational model’s accuracy

###### Round 2 - Learning-Informed Design

– Integration of SPR results from Round 1, learning from 11 expression failures and 85+ designs showing >10³ nM binding
– Generation of project-specific model weights enhancing ability to predict high-affinity binders
– The direct learning head architecture maintains complete separation: foundational model parameters remain frozen while project-specific weights adapt to experimental data, ensuring proprietary information remains isolated and secure
– Re-evaluation of original 6 × 10⁸ compendium using updated project-specific model
– Selection of 96 new designs with recalibrated confidence scores informed by Round 1 outcomes

###### Round 3 - Convergence and Validation

– Selection of 96 candidates incorporating all learned constraints from previous rounds:

– CDR-H3 length: 10-15 amino acids with appropriate flexibility motifs
– Hydrophobicity: GRAVY score −0.2 to 0.3 for optimal folding
– Systematic avoidance of hydrophobic patches in CDR regions identified as problematic
– **SPR Validation Results:**

– 20 candidates achieved KD ≤100 nM (21% success rate)
– 3 candidates demonstrated sub-10 nM affinities
– Best candidate: 0.84 nM binding affinity
– Substantial improvement from Round 1’s <10% success rate

**Figure 4:**
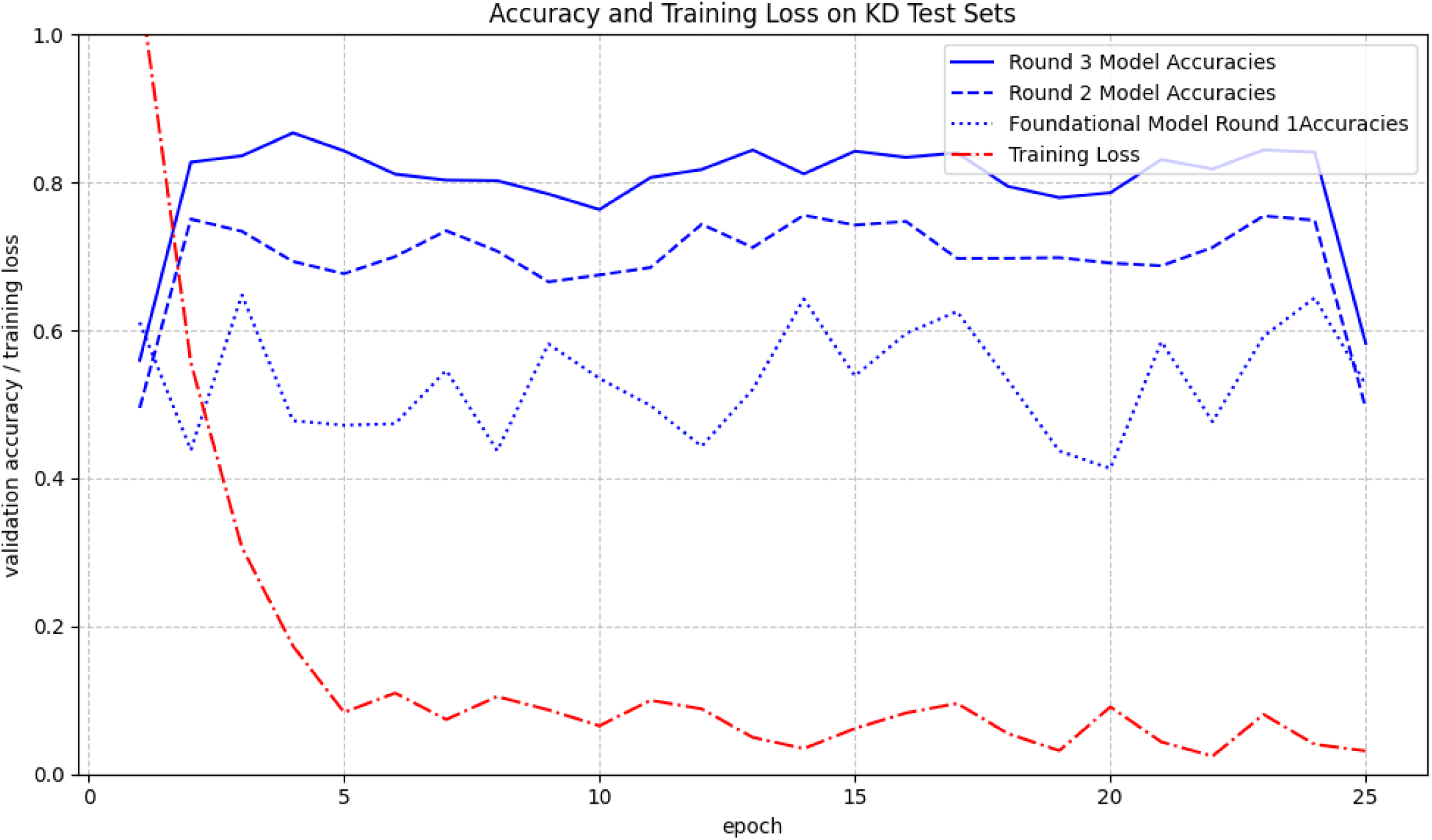
Model Training Performance Across Iterative Rounds. Training accuracy and loss metrics demonstrating iterative model improvement through experimental feedback integration. The foundational model (dotted blue line) shows baseline performance, while Round 2 (dashed blue line) and Round 3 (solid blue line) models demonstrate progressive accuracy improvements from 65% to 85% through project-specific direct learning head model training. Training loss (red dashed line) decreases consistently across iterations, indicating effective learning from experimental SPR data. Each round of experimental feedback enables the model to better discriminate between binders and non-binders for the specific chemokine receptor target.

#### 3.3.4 Experimental Validation and Comprehensive Results

##### Expression and Purification Protocol

All selected scFv panel sequences underwent systematic experimental validation:

– Gene synthesis as gBlocks Gene Fragments (Integrated DNA Technologies) and cloning into pTT5 expression vector (National Research Council Canada)
– Transient transfection into HEK293-6E cells using PEI MAX (Polysciences) at a 1:3 DNA:PEI ratio
– Expression at 37°C with 8% CO2 in FreeStyle 293 Expression Medium for 7 days
– Purification using protein A chromatography (MabSelect SuRe, Cytiva) followed by size exclusion chromatography
– Verification of >95% purity by SDS-PAGE under both reducing and non-reducing conditions [113]

##### Biophysical Characterization

– **Size-exclusion chromatography (SEC)** [114]: Round 1 showed 88% of constructs displayed predominant monomeric peaks with retention volumes consistent with high monomeric content (∼12 mL), while 11 sequences showed inclusion body formation or aggregation. By Round 3, expression success reached nearly 100%.
– **Aggregation assessment** [115]: Several variants exhibited delayed retention times, indicating mild aggregation or stickiness under assay conditions but still within acceptable ranges

##### Binding Kinetics and Affinity Measurements

Binding kinetics were comprehensively measured using surface plasmon resonance (SPR) on Biacore 8K (Cytiva) against the chemokine receptor extracellular domain. Each scFv was captured via anti-His antibody followed by injection of a receptor at seven concentrations (0.78-200 nM) to determine association and dissociation rate constants (Kon and Koff) for equilibrium dissociation constant calculation [116]. Refer to Figure 6 for several sensorgrams resulting from this project.

##### Progressive Results Across Iterative Rounds

###### Round 1

– Despite representing the highest computational confidence predictions from 6 × 10⁸ candidates, majority of antibodies exhibited poor binding affinities (KD > 10³ nM)
– High prevalence of non-specific binding attributed to unreliable curve fitting during SPR analysis
– Critical learning: High computational confidence did not translate to experimental success without incorporating actual failure data

###### Round 2

– Significant improvements observed with several candidates achieving KD values between 10² and 10³ nM
– 3 candidates achieved sub-100 nM binding
– Clear evidence of model learning from experimental feedback integration
– Definition of “high confidence” fundamentally recalibrated based on Round 1 outcomes

###### Round 3

– Marked enrichment of strong binders with majority achieving KD values below 10² nM
– **20 candidates demonstrated KD ≤ 100 nM, representing a 21% success rate**
– **3 candidates achieved sub-10 nM KD affinities, with the best candidate reaching 0.84 nM**
– Substantial reduction in non-specific binding and low-concentration artifacts
– Validation that learning from comprehensive failure characterization enables dramatic performance improvements

**Figure 5:**
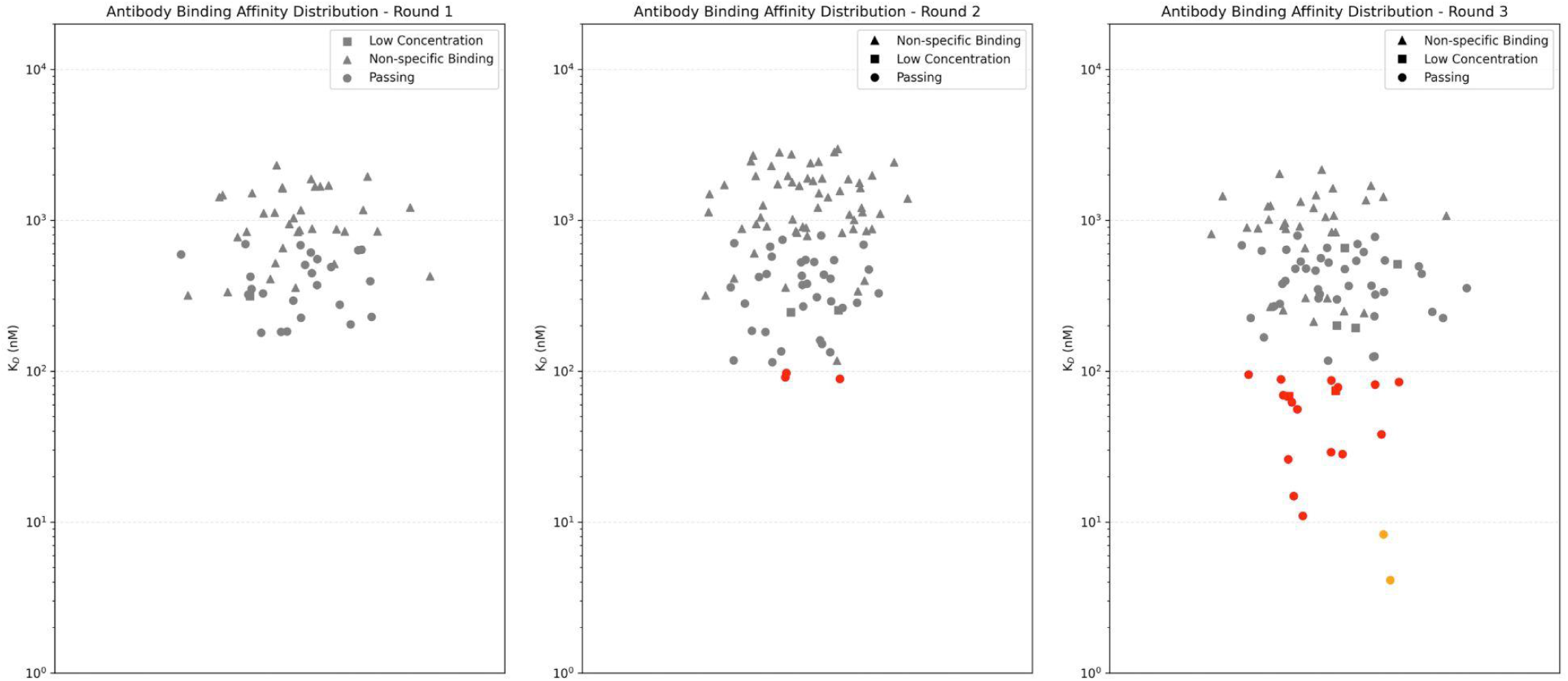
Antibody Binding Affinities, Round 1, Round 2, and Round 3. Binding affinity distributions for antibodies across three experimental rounds, with KD values plotted on a logarithmic scale. Antibodies are categorized as **Passing**, **Non-specific Binding**, or **Low Concentration**. • **Round 1:** A majority of antibodies exhibited KD values above 10^3 nM, indicative of poor binding affinity despite being selected as the highest computational confidence predictions. Non-specific binding in this range is attributed to unreliable curve fitting during SPR analysis. Low-concentration antibodies also contributed to the scarcity of strong binders. • **Round 2:** Improvements are observed, with a subset of antibodies achieving KD values below 10^3 nM. However, non-specific binding remains prevalent among antibodies with high KD values, reflecting challenges in reliable binding characterization. • **Round 3:** A marked enrichment of strong binders is observed, with the majority of antibodies achieving KD values below 10^2 nM. The reduction in non-specific binding and low-concentration antibodies indicates enhanced selection and improved design across iterations. The achievement of 0.84 nM binding affinity validates the power of learning from comprehensive failure characterization. This progression underscores the iterative optimization of antibody candidates, addressing issues of non-specific binding and low-concentration effects to identify high-affinity binders.

**Figure 6:**
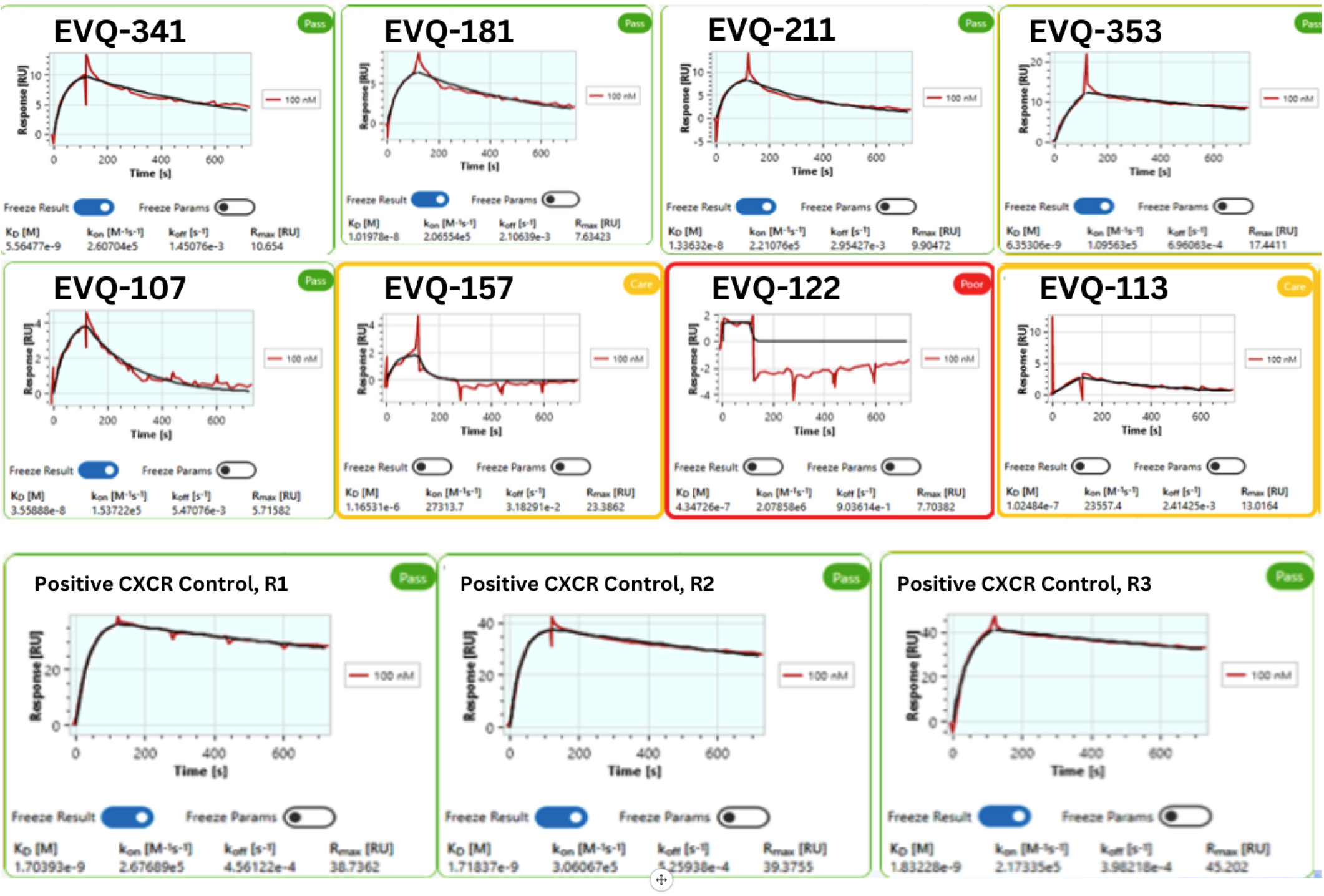
Representative SPR sensograms validating binding affinity data from antibody candidates across experimental rounds. Sensograms of selected antibody candidates (top and middle row), demonstrating moderate to high binding profiles with KD values approaching the passing threshold. The last three (EVQ-157, EVQ-122, EVQ-113) depicted poor to no binding. A control antibody used as a benchmark, displaying expected binding behavior and reliable curve rounding in each of the three rounds (bottom row). The progressive improvement in binding characteristics from Rounds 2 to 3 highlights the success of iterative optimization and selection strategies, with the control antibody providing validation of the experimental setup and SPR analysis.

#### 3.3.5 Project Timeline and Resource Efficiency

##### Total Project Duration

6 months from initial model training to final optimized candidate panel delivery

##### Rapid Experimental Turnaround

Each round achieved remarkably fast timelines—just 6 weeks from project initiation to SPR assay results, demonstrating the efficiency of the multi-shot learning approach:

– Week 1-2: Computational screening of 6×10⁸ candidates and selection of 96 designs
– Week 3-4: Gene synthesis and cloning
– Week 5: Expression and purification
– Week 6: SPR characterization and data analysis

##### Detailed Phase Breakdown

– **Months 1-2**: Initial Hyperbind2 model training, sequence generation, and first-round SPR validation with 96 candidates (6 weeks active, 2 weeks analysis/planning)
– **Months 3-4**: Experimental result integration, model refinement, and second-round sequence generation and validation
– **Months 5-6**: Final optimization round, experimental evaluation, and delivery of optimized candidate panel

This compressed timeline—from computational design to experimental results in 6 weeks per round—contrasts sharply with traditional approaches requiring 3-6 months per screening round. The rapid iteration enabled by our multi-shot learning architecture allows three complete design-build-test cycles in the time traditional methods require for a single round.

##### Resource Efficiency Demonstration

The computational approach enabled remarkable efficiency gains:

– Computational evaluation of 6 × 10⁸ potential candidates
– Experimental synthesis and testing of only 288 candidates total (96 per round)
– 4-5 orders of magnitude reduction in experimental screening burden compared to traditional library approaches
– Rapid identification of multiple sub-10 nM binders within 6-month timeline, including best-in-class 0.84 nM binder
– Comprehensive learning from all experimental outcomes, including failures that would conventionally be discarded

## 4. Contextualizating Success: Fundamental Constraints in Computational Antibody Design

### Current Limitations and Active Research Directions

Despite Hyperbind2’s demonstrated success, we acknowledge specific limitations that define ongoing research priorities:

### Epitope Uncertainty and Validation

While our approach successfully identifies high-affinity binders, confirming that antibodies bind to intended epitopes rather than alternative sites remains a challenge inherent to sequence-based methods. The 20 sub-100nM binders identified in our chemokine receptor campaign require additional characterization to confirm epitope specificity. Current internal research addresses this through:

– **Digital epitope mapping**: Computational alanine scanning across the antigen surface to predict critical binding residues
– **Ablation studies**: Systematic testing of antigen variants with mutated epitope regions
– **Competitive binding assays**: Validation that computationally designed antibodies compete with known epitope-specific binders
– **Integration with structural methods**: When binders are identified, employ cryo-EM or crystallography for definitive epitope confirmation

As noted in recent work from Chai Discovery [117], “further competitive binding assays and laboratory 3D structure determination should be performed” to confirm epitope targeting. We concur with this assessment—while computational design can achieve impressive hit rates, epitope specificity requires additional experimental validation. Our current research focuses on developing confidence scores that predict not just binding likelihood but also epitope targeting probability, enabling prioritization of candidates most likely to bind at therapeutically relevant sites.

### Reduced Mechanistic Explainability

The sequence-only inference approach, while computationally efficient, provides limited mechanistic insight into why specific antibodies bind. Unlike structure-based methods that can visualize predicted binding interfaces, our embeddings operate in high-dimensional spaces that resist low-dimensional interpretation.

### Functional Activity Prediction

Current optimization focuses primarily on binding affinity, which doesn’t always correlate with functional activity. Distinguishing agonists from antagonists, predicting neutralization potency, or identifying allosteric modulators requires additional experimental characterization beyond SPR binding data.

These limitations represent active research areas rather than fundamental barriers. The multi-shot learning framework naturally accommodates additional validation data—epitope mapping results, competitive binding assays, and functional characterization can all be incorporated as training signals in subsequent rounds, progressively improving both affinity and specificity predictions.

### 4.1 The Persistent Reality of High Failure Rates

Despite processing millions of training sequences and achieving progressive improvements across multiple campaigns—including our chemokine receptor (20 sub-100nM binders), DPP4 (successful VHH-Fc binders), intracellular protease targets, and immunomodulatory receptors—computational antibody design across the field faces fundamental reliability challenges that vary dramatically with target characteristics. All current computational approaches encounter scenarios where results fall far short of desired thresholds—ranging from panels with only weak or moderate binders to complete failures where no candidates meet the empirical criteria. This target-dependent variability reflects the diverse phenotypes that proteins present for antibody recognition: some targets offer multiple immunogenic epitopes with favorable binding geometries, while others present cryptic, conformationally unstable, or poorly accessible surfaces that resist antibody engagement regardless of whether the discovery approach is in silico, in vivo, or in vitro.

The field’s collective hope that more data would solve prediction accuracy has proven optimistic; despite having more sequence-function pairs than a decade ago, challenging targets often yield sub-optimal results with binding affinities orders of magnitude weaker than therapeutic requirements, while favorable targets may reach acceptable success rates of 40-50%. This variability isn’t a failure of any particular approach but reflects the inherent complexity of protein-protein recognition across diverse biological contexts.

### 4.2 The Measurement Problem: Inconsistent Ground Truth Across Assays

A fundamental challenge that no amount of machine learning sophistication can overcome is the inconsistency of experimental measurements themselves. An antibody showing sub-nanomolar affinity in SPR may show no binding in ELISA, weak signal in flow cytometry, and complete inactivity in cell-based functional assays. These aren’t measurement errors but reflect different aspects of molecular recognition under different conditions—immobilized versus solution-phase antigen, monomeric versus multimeric presentation, presence versus absence of cellular cofactors. Models trained on SPR data learn to predict SPR outcomes, not biological reality. This creates an epistemological crisis: we’re optimizing for proxy measurements that may have minimal correlation with therapeutic efficacy. The notion of a single “ground truth” for antibody-antigen interaction is a convenient fiction that obscures the multidimensional nature of molecular recognition.

### 4.3 Systematic Library Failures and Unknown Unknowns

In some cases, even sophisticated diversity-optimized libraries designed to broadly sample sequence space while maintaining structural integrity fail to yield binders significantly above background in FACS sorting. When these failures occur, they suggest that current computational approaches may be missing critical determinants of binding for certain target classes. For these challenging targets, we may be exploring suboptimal dimensions of sequence space, focusing on parameters that correlate weakly with actual binding while missing unknown factors that dominate recognition. However, in other campaigns, the same computational strategies yield rich panels of functional binders, indicating that our platform’s current understanding—hydrophobicity patterns, charge complementarity, shape matching—works well for certain protein classes while being insufficient for others. This target-dependent success pattern implies that universal rules for antibody-antigen recognition remain elusive, with each target class potentially requiring distinct optimization strategies.

### 4.4 Experimental Noise as an Absolute Ceiling

The theoretical limit of any predictive model is bounded by the reproducibility of its training data, and antibody assays are notoriously irreproducible. Antigens from different expression batches can show 10-fold affinity variations due to glycosylation differences or misfolding. SPR chips degrade over time, introducing systematic drift. Temperature fluctuations of 2°C can shift KD values by an order of magnitude. Reagent lot changes can invert rank ordering of candidates. Even technical replicates within the same experiment routinely show coefficients of variation exceeding 30%. When the experimental noise approaches or exceeds the biological signal, no amount of computational sophistication can extract reliable patterns.

### 4.5 The Nonlinear, Chaotic Nature of Protein Interactions

Perhaps most fundamentally, antibody-antigen interactions exhibit chaotic sensitivity to initial conditions that defies systematic prediction. Single point mutations can eliminate binding entirely or improve affinity 1000-fold, with no reliable way to predict which outcome will occur. Enthalpy-entropy compensation means that improving one aspect of binding often worsens another in unpredictable ways. Allostery and induced fit create context-dependent conformational landscapes where the act of binding changes the target. Water molecules and ions at the interface contribute binding energy comparable to direct protein contacts but rearrange on nanosecond timescales beyond current modeling capabilities. We’re not dealing simply with a complicated problem—we’re confronting irreducible biological complexity where deterministic prediction may be theoretically impossible.

### 4.6 Operating Within an Imperfect Reality

These limitations aren’t unique to Hyperbind2 or even computational approaches broadly—they reflect fundamental constraints of biologics discovery that all methods face. High-throughput screening campaigns with million-member libraries sometimes yield no hits despite extensive effort. Even animal based immunization, the oldest of discovery approaches, yields variable success rates. Lead optimization campaigns can plateau at local maxima regardless of resources invested. All current methods—computational and experimental—operate within biological systems where partial patterns exist but complete prediction remains elusive.

Given this reality, computational design should serve as a complementary track alongside traditional experimental approaches, not as a replacement. Direct learning with iterative lab-to-AI feedback enhances rather than supplants established discovery workflows. While library screening explores vast sequence diversity through physical selection, computational methods can investigate orthogonal regions of sequence space informed by different principles. When experimental screening identifies initial hits, computational modeling can guide selection and optimization. This bidirectional information flow between computational and experimental tracks strengthens both approaches.

The economics now make dual-track strategies compelling: computational screening costs a fraction of wet-lab validation, making it prudent to include digital design as a supplementary exploration method in discovery campaigns. Generating and ranking 10^8 computational designs adds relatively low cost while potentially identifying sequences that complement library-derived candidates. The improvements reduce the number of variants from 10^8 to 10^5, improve success rates through iterative refinement, ultimately accelerating optimization cycles that enhance rather than replace traditional discovery methods.

This integration of computational prediction with experimental validation, where each informs and improves the other through multi-shot learning cycles, represents a paradigm shift. Not computation replacing experimentation, but both working in concert, extracting maximum value from every experimental outcome to progressively refine our understanding of what antibodies.

## 5. Iterative Learning Compared to Single-Pass Prediction Methods

### 5.1 Comparative Benchmarking with Leading Platforms

A comparative analysis of HyperBind2, RoseTTAFold-Diffusion (RF-D), Nabla Bio’s JAM, and Chai Discovery’s Chai-2 was performed to position each within the broader landscape of computational antibody design [46, 118, 119]. Each framework reflects a distinct philosophy: HyperBind2 emphasizes sequence-only antibody-antigen generalizability with epitope granularity and learning from experimental failures; RF-D advances atomic-level backbone generation; JAM focuses on epitope-anchored complex design, and Chai-2 pursues integrative antibody–antigen co-design with a focus on structure-guided optimization.

#### Note on comparability

The results below are presented as reported in their respective manuscripts. Because assay formats, hit-rate definitions, and validation pipelines differ, values should not be interpreted as a direct head-to-head benchmark.

**Table 1.** Comparison of leading computational antibody design platforms.

| Dimension | HyperBind2 | RoseTTAFold-Diffusion (RF-D) | Nabla JAM | Chai-2 |
| --- | --- | --- | --- | --- |
| Core model type | Contrastive pre-trained language model (sequence → embedding, ranking) over pre-screened human sequence library | Diffusion on 3D residue frames; sequence via ProteinMPNN. | Generative model over antibody–antigen complexes | All-atom structure-conditioned generative model with integrated folding; designs binders conditioned on a binding site specified by a few epitope residues. |
| Primary output | Ranked variable-domain antibody sequences (scFv, VHH, IgG) | De novo backbones + sequences | Full antibody–antigen complex design (sequence + structure) | Antibody–antigen pair designs (scFv, VHH, miniproteins) with epitope-specified conditioning. |
| Input requirements | Antigen sequence, (structure optional; epitope/region optional) | Unconditional or structure-conditioned: symmetry/motif scaffolding, or binder design given a target structure + specified interface hotspots. | Target sequence and/or structure plus epitope residues; structure used as a flexible constraint. | Target protein structure plus up to four epitope residues used as a prompt. |
| **Reported hit-rate** | 7.9% to 16.7% experimental VH/VL binder success (<100nM); top binders with <10nM KD; all experimental designs with sequence-based target information data across multiple targets | 19% experimental binder success (BLI) across five targets; nanomolar KDs; cryo-EM complex matches design (RMSD ~0.63 Å). Fold-specified in-silico success ~42–54%. | ~0.5% at 100 nM for a PDB-absent antigen, with top 12–24 nM KD after recombinant characterization; Paired scFv combinatorial library yielded ~0.0017% bind rate with double-digit nM mAb affinities. | ~15.5% average experimental hit rate overall (20.0% VHH; 13.7% scFv) with ≥1 binder for ~50% of targets; epitope localization to be experimentally confirmed. |
| **Design runtime** | >1 million sequences per hour; ~10<sup>8</sup> ranked in <48 h. | ~11 s per 100 aa design on RTX A4000; inference can be accelerated by fewer timesteps with high in-silico success; AF2 filtering used for selection. | 4–6 weeks from design to recombinant characterization. | In-silico pipeline ~1 day; wet-lab validation ~2 weeks (single round reported); no per-design GPU time stated. |
| **In-vitro strategy** | Multi-round recombinant antibody expression with SPR validation | Expression/purification → BLI screening/titrations; cryo-EM complex of a designed binder; AF2 used for in-silico validation/filtering. | Two-stage: yeast display (MACS where applicable + two FACS rounds, NGS) → recombinant production (VHH-Fc or mAb), BLI/on-cell binding, and developability. | Single-round testing (≤20 designs/target); BLI hit-calling with predefined thresholds; additional specificity/developability assays; epitope mapping planned. |
| **Library scale** | Compendium of 108–109 sequences screened; 96 expressed per round (288 total tested) | ~95 designs per target taken forward for lab testing after AF2 filtering (paper does not claim 10<sup>5</sup>-scale batches). | Typically ~10<sup>4</sup> designs/target (pooled across targets in some screens); scFv combinatorial ~9×10<sup>6</sup>; empirical neighborhood exploration up to ~4×10<sup>8</sup> variants. | ≤20 designs per target taken forward to the lab. |
| **Notable strengths** | Functions without structure; Offers epitope-specific, developable designs; scalable across hard/undruggable targets | Unconditional and conditioned generation; fold control (e.g., TIM/NTF2), atomic-level agreement in a binder–target cryo-EM complex. | Epitope-specific designs with favorable developability | Programmable epitope-specified design; multiple modalities; cross-reactivity steering; double-digit experimental hit rates across >50 novel targets. |
| **Access model** | Open-source version described, available on Github; commercial platform designs via fee-for-service | Open-source code released; see data & code availability at article DOI | Proprietary (Nabla Bio platform); access via partnerships | Company platform (access model not specified in manuscript). Given recent \$70M funding, likely access via partnerships |

#### Narrative Comparison

##### Technical differences

HyperBind2 is unique in that it requires only an antigen sequence, enabling the design of antibodies against targets lacking solved structures. RF-Diffusion generates backbones by denoising residue-frame coordinates and supports unconditional generation as well as conditioned tasks; sequences are assigned with ProteinMPNN. JAM generatively designs antibody–antigen complexes conditioned on target sequence and/or structure with user-specified epitope residues, treating structure as a flexible constraint rather than a hard requirement. Chai-2 generates binders conditioned on a defined target structure and a binding site specified by up to four epitope residues; modalities include scFv, VHH, and miniproteins.

##### Experimental strategy

HyperBind2 emphasizes iterative multi-round design–build–test, with 96 recombinant antibodies expressed per round (288 total) and relies on third-party/client-driven production and testing; 20 binders were ≤100 nM, including 3 sub-10 nM, validated by SPR. For RF-Diffusion, across five targets, 19% of designs showed binding by BLI (≥50% of positive control), including nanomolar affinities; one designed binder–target complex was determined by cryo-EM and matched the design at ∼0.63 Å RMSD. JAM uses a two-stage workflow: MACS/FACS-based yeast display (typically two FACS rounds) followed by recombinant characterization (BLI or on-cell) and developability assays with an overall 4–6-week design-to-characterization timeline. Chai-2 uses one round of testing (≤20 designs per target) with BLI-based hit calling (signal >0.1 nm and >300% of background), followed by targeted characterization; no iterative rounds were performed in this study.

##### Throughput and scalability

HyperBind2 can screen >1 million sequences per hour and rank ∼10^8 designs (from a pre-screened, fully human library) in under two days, supporting rapid iteration across multiple antigens; 96 are expressed per round for testing. RF-Diffusion generates a 100-aa design in ∼11 s on an RTX A4000, with designs filtered by AF2 confidence. Typical JAM designed libraries are ∼10^4 per target (pooled across targets in some screens), with scFv combinatorial libraries at ∼9×10^6 and optional empirical neighborhood exploration reaching ∼4×10^8 variants in a single design round. Chai-2 emphasizes quality-controlled, small-batch evaluation (≤20 designs/target) rather than high-throughput libraries; the in-silico stage is ∼1 day and wet-lab ∼2 weeks.

#### Platform Technology Access

##### HyperBind2 (EVQLV)

HyperBind2’s defining strength is its ability to operate without structural inputs, making it particularly valuable for antigens where no solved structure exists or where structural models remain uncertain. The platform combines an open-source sequence-based model for computational scientists with a commercial fee-for-service offering (via abtique.com) that enables antibody scientists in biopharma, startups, and academia to access AI-driven antibody design without specialized training or infrastructure. This dual-access model positions HyperBind2 as a democratized platform, lowering the barrier for immediate experimental integration while still supporting computational exploration at scale.

##### RF-Diffusion (University of Washington, Xaira Therapeutics)

RF-Diffusion provides a powerful route to de novo backbone generation and scaffold innovation with atomic precision, validated by both novel folds and binder-target cryo-EM complexes. Access is available through its open-source codebase, though practical use requires significant computational expertise, coding skills, and downstream validation infrastructure. Beyond academia, the platform’s commercial trajectory is being advanced through Xaira Therapeutics, the company formed from the RF-D work. As a result, RF-D is most suited to computational biologists and structural biologists with coding capacity that are seeking fine-grained backbone design.

##### JAM (Nabla Bio)

JAM’s strategic role is in generating antibody–antigen complexes anchored on precise epitope definitions, followed by rapid experimental confirmation and developability profiling. Unlike HyperBind2 or RF-D, JAM is not broadly available: access appears restricted to proprietary use within Nabla Bio and through large-scale biopharma partnerships. Recent collaborations with Bristol Myers Squibb, Takeda, and AstraZeneca underscore its positioning for targeted campaigns through partnerships for enterprise-level discovery.

##### Chai-2 (Chai Discovery)

Chai-2 advances antibody–antigen co-design with explicit epitope prompting, supporting modalities such as scFv, VHH, and miniproteins. It demonstrates double-digit experimental hit rates across >50 novel targets, though the precise epitope fidelity of those binders remains to be experimentally confirmed. Access today is partnership-driven, making it most relevant to biopharma collaborations seeking integrative antibody design. Its strategic appeal lies in coupling next-generation AI design with translational partnerships at scale.

**Table 2.** Recommended platform selection by discovery scenario.

| Use case | Recommended Model | Rationale | Access model | Primary users |
| --- | --- | --- | --- | --- |
| Limited structural information; rapid in silico-to-lab screen | HyperBind2 | Sequence-only input enables fast iteration without 3D antigen structures | Open-source, research-only license; commercial access through fee-for-service (via abtique.com) | Antibody discovery and engineering scientists in biopharma, startups, and academia; computational biologists |
| Novel scaffold discovery, atomic-precision folds | RF-D | Diffusion model enables backbone generation with fine structural control | Open-source license | Computational biologists and structural biologists with coding skills |
| Epitope-specific, structure-anchored design | JAM | Complex modeling supports atomic placement at known binding sites | Proprietary platform; access through partnerships | Biopharma discovery via partnering |
| Epitope-specified paired designs | Chai-2 | Co-design conditioned on user-defined epitope residues; experimental localization remains to be confirmed | Proprietary platform; access through partnerships | Biopharma discovery via partnering |

## Conclusion

Together, these platforms illustrate the rapidly diversifying landscape of computational antibody design. The comparative benchmarking highlights that each platform advances antibody design through a distinct philosophy and access model, serving complementary roles in the therapeutic discovery ecosystem. Collectively, these approaches are not mutually exclusive but rather complementary: HyperBind2 lowers the barrier for broad experimental adoption of AI-assisted antibody design; RF-Diffusion expands the boundaries of structural innovation; JAM anchors discovery in precise epitope targeting for large pharma campaigns; Chai-2 advances epitope-specified co-design through strategic collaborations. Together, they chart a multifaceted path for how AI will reshape antibody discovery and development, offering distinct yet synergistic strengths across access, scalability, and translational impact.

### 5.2 Platform Scalability and Additional Validation

#### Multi-Format Therapeutic Support

Hyperbind2 has been validated across diverse therapeutic formats beyond the primary case study:

– **Single-domain antibodies (VHHs)**: Successful identification of high-affinity nanobodies with favorable biophysical properties
– **scFv formats**: Demonstrated comprehensively in the chemokine receptor case study
– **Full-length IgG antibodies**: Validation for conventional therapeutic formats with proper assembly and function
– **Complex therapeutic formats**: Preliminary validation for next-generation formats including CARs, BiTEs, and bispecific antibodies

#### Additional Case Study Results

Platform effectiveness has been demonstrated across multiple challenging targets:

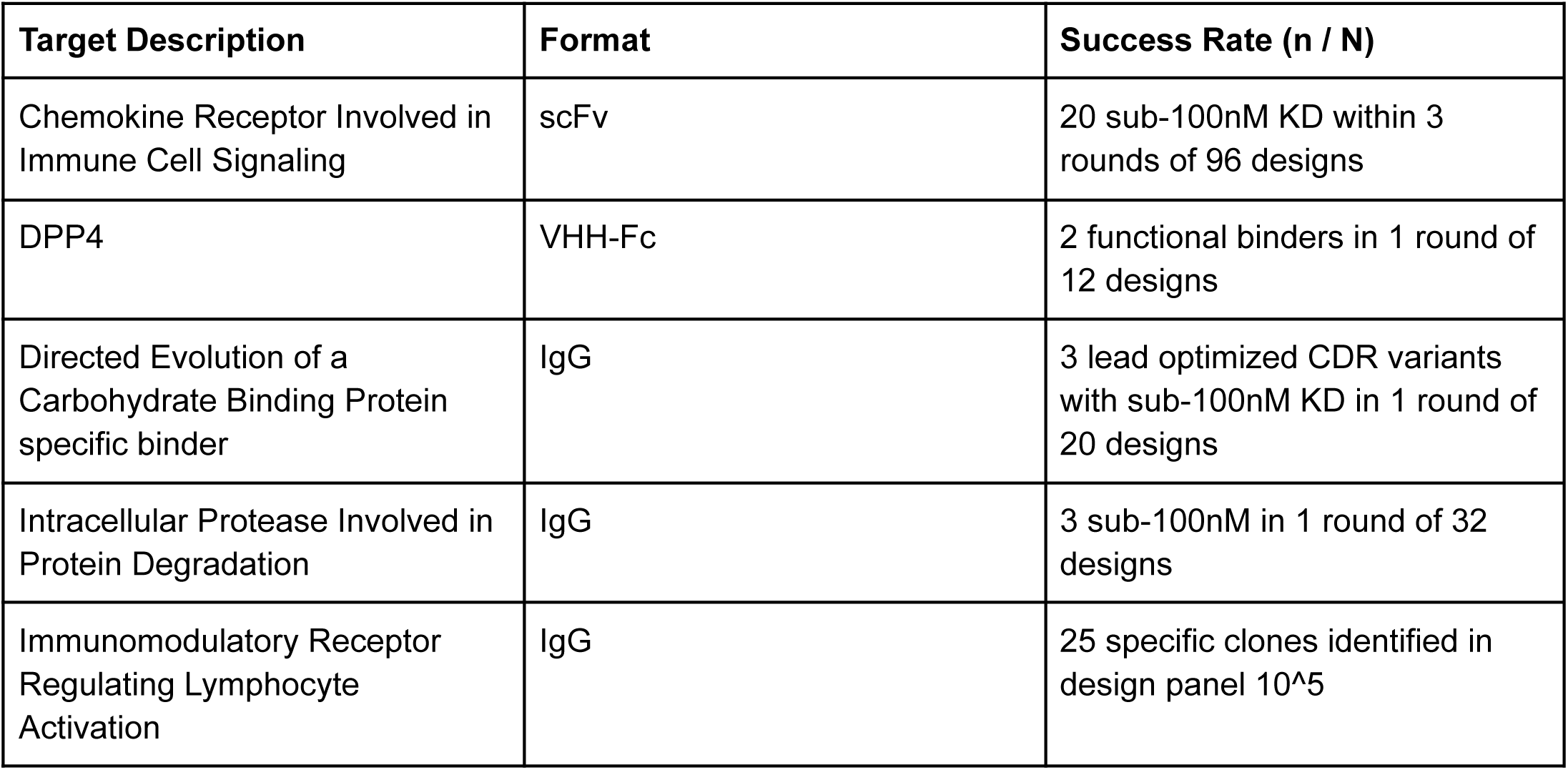

| Target Description | Format | Success Rate (n / N) |
| --- | --- | --- |
| Chemokine Receptor Involved in Immune Cell Signaling | scFv | 20 sub-100nM KD within 3 rounds of 96 designs |
| DPP4 | VHH-Fc | 2 functional binders in 1 round of 12 designs |
| Directed Evolution of a Carbohydrate Binding Protein specific binder | IgG | 3 lead optimized CDR variants with sub-100nM KD in 1 round of 20 designs |
| Intracellular Protease Involved in Protein Degradation | IgG | 3 sub-100nM in 1 round of 32 designs |
| Immunomodulatory Receptor Regulating Lymphocyte Activation | IgG | 25 specific clones identified in design panel 10<sup>5</sup> |

These results demonstrate consistent performance across diverse target classes, antibody formats, and screening scales.

## 6. Discussion

### 6.1 A Paradigm Shift in Antibody Discovery

The success demonstrated in our chemokine receptor case study—achieving 20 sub-100nM binders including a best-in-class 0.84 nM affinity with just 288 tested sequences—represents a fundamental paradigm shift in therapeutic antibody discovery. Where traditional campaigns require screening 10^10-10^12 variants across 8-12 experimental rounds, and where the best reported traditional campaign for this target achieved only a single 250 nM binder after massive library screening, our multi-shot learning approach achieved a 21% success rate in just 3 rounds.

This transformation addresses a critical gap in current computational antibody design: the inability to efficiently learn from experimental feedback. Zero-shot methods like CHAI-2 achieve impressive ∼16% initial hit rates but remain static—their billion-parameter generative architectures were designed for large-scale pre-training, not adaptation from small experimental datasets. While these models could theoretically incorporate experimental data, fine-tuning faces insurmountable barriers: determining optimal stopping points with limited validation data risks catastrophic forgetting, computational costs reach hundreds of GPU-hours per project, and critically, pharmaceutical projects cannot accept the IP risks of data entanglement in shared model weights. The alternative—maintaining separate billion-parameter models per project—multiplies already prohibitive costs.

Our multi-shot learning architecture solves this through purpose-built design for iterative adaptation. Through synthetic amplification, 96 experimental tests generate approximately 9,000 training pairs via biologically-informed pairing strategies that extract maximum learning from each observation. This enables robust model adaptation while maintaining complete data segregation between projects. Unlike generative models that must balance broad capabilities against project-specific learning, our contrastive framework naturally incorporates experimental evidence without disrupting foundational representations. The system updates within 24-48 hours, enabling true iterative optimization where Round N directly informs Round N+1 predictions.

This architectural advantage manifests in progressive performance improvements. Starting from computational predictions yielding only 5-10 weak positives (100-1000nM) in Round 1, the system learned from these outcomes to achieve 3 sub-100nM binders in Round 2, then refined further to achieve 20 sub-100nM binders in Round 3. This progression—from <10% to 21% success rate—demonstrates the power of multi-shot learning versus static prediction, achieving in 3 rounds what traditional approaches fail to achieve in 8-12 rounds.

This multi-shot capability enables compound learning effects where each round builds on accumulated knowledge. Round 1 explores broadly across sequence space, identifying initial binding signals and learning which sequence features correlate with even weak activity. Round 2 applies these learned patterns to focus on promising regions while avoiding confirmed failure modes. Round 3 further refines based on accumulated insights, converging on optimal solutions. This iterative refinement transforms antibody discovery from independent screening rounds to a unified optimization trajectory where knowledge compounds rather than resets.

The implications reshape therapeutic development. Small biotechs and academic groups can now pursue challenging targets previously requiring big pharma resources. The ability to start with computational predictions and rapidly improve through experimental feedback democratizes access to AI-driven discovery. Projects can adapt to target-specific challenges—expression systems, assay conditions, developability requirements—rather than hoping pre-trained models generalize appropriately. Most importantly, the approach enables systematic exploration of previously intractable targets like the multi-pass membrane protein in our study, where structural methods cannot be applied and traditional screening has repeatedly failed.

This represents more than technical advancement—it’s a fundamental shift in how we approach antibody discovery. Instead of massive parallel screening hoping for lucky hits, or static computational predictions that cannot improve, we enable true human-AI collaboration where computational predictions and experimental validation create a virtuous cycle of continuous improvement. Each experiment teaches the system, each round gets smarter, and challenging targets become progressively more tractable. This transforms antibody discovery from a game of chance to a systematic optimization process.

### 6.2 Theoretical Foundation of Multi-Shot Learning

The multi-shot learning capability demonstrated in our chemokine receptor study rests on four theoretical pillars that together enable efficient learning from limited experimental observations.

Sample Efficiency Through Transfer Learning: Our frozen foundation model provides rich protein representations learned from millions of sequences, capturing evolutionary patterns and structural constraints. The innovation lies in the lightweight adaptation layer that requires only 10-100 experimental examples to map these representations to user-specific outcomes. This transfer learning approach preserves general protein knowledge while enabling rapid specialization—analogous to how a seasoned scientist applies broad expertise to a new specific problem.

Contrastive Learning Amplification: Each experimental batch undergoes synthetic amplification through systematic pairing strategies. From 96 experiments, we generate approximately 9,000 training pairs through biologically-informed contrasts: sequences with similar features but different outcomes, systematic mutations around decision boundaries, and cross-family comparisons. This O(n²) scaling transforms limited observations into comprehensive training signals, enabling the model to learn nuanced distinctions between success and failure.

Bayesian Posterior Refinement: Each round updates the posterior distribution over sequence space in a principled Bayesian framework. Round 1 begins with broad, uninformed priors derived from the foundation model. As experimental data accumulates, these priors sharpen into focused posteriors concentrated on high-probability binding regions. This mathematical framework explains the progressive improvement—the model isn’t just memorizing successful sequences but building probabilistic models of what makes sequences successful for this specific target.

Active Learning Principles: Our selection strategy for each round follows active learning theory, choosing sequences that maximally reduce model uncertainty. Rather than random sampling or pure exploitation of predicted binders, we balance exploration of uncertain regions with exploitation of promising areas. This uncertainty-guided selection accelerates convergence, achieving in 3-4 rounds what random sampling would require 10+ rounds to accomplish.

These theoretical foundations translate directly to practical advantages: 10-100x fewer experiments needed, 24-48 hour learning cycles instead of weeks, and progressive improvement rather than static predictions. The theory isn’t abstract—it’s validated by the 0.84 nM binder discovered through exactly these principles.

### 6.3 Advantages of Embedding-Driven Architecture

The embedding-driven architecture employed by Hyperbind2 offers critical advantages that enable practical multi-shot learning where other approaches fail.

Computational Efficiency at Scale: Our embedding framework processes 10⁸-10⁹ candidate sequences in under 48 hours using standard GPU infrastructure—orders of magnitude faster than structure-based methods requiring molecular dynamics or docking simulations. This speed is essential for iterative optimization; if each round required weeks of computation, the feedback loop would be too slow for practical drug discovery timelines.

Natural Compositionality with Experimental Data: Embeddings provide a continuous representation space where experimental outcomes can be directly incorporated. Unlike generative models that must balance preserving their generative capabilities with incorporating new information, or classification models that require redefining decision boundaries, our contrastive embeddings naturally adapt to new positive-negative pairs without architectural modifications. This compositionality means adding Round 2 experimental data doesn’t disrupt Round 1 learning—knowledge accumulates rather than overwrites.

Target-Agnostic Transfer: The embedding space learned from diverse antibody-antigen pairs transfers effectively to new targets without target-specific architectural changes. This differs from structure-based approaches requiring target-specific force fields or epitope-anchored methods needing predefined binding sites. The same embedding architecture that succeeded with our chemokine receptor has been validated across membrane proteins, enzymes, and cytokines without modification.

Interpretable Attention Patterns: While maintaining computational efficiency, our attention mechanisms provide interpretability into learned binding patterns. Analysis of attention weights revealed the model discovering known binding motifs (RYY, YWW, GSS flexibility patterns) without explicit programming. This interpretability builds trust with experimental teams and provides hypotheses for rational design improvements.

### 6.4 Significance of Rapid Iterative Feedback Mechanism

The 24-48 hour feedback loop between experimental validation and model updating fundamentally changes how antibody discovery campaigns operate. Traditional computational approaches operate in batch mode—generate predictions, test extensively, analyze results over weeks or months, then start fresh with new predictions. Our rapid iterative mechanism transforms this into a continuous optimization process where Monday’s SPR results improve Tuesday’s predictions, creating unprecedented momentum in discovery campaigns. This speed is enabled by architectural decisions prioritizing adaptation efficiency. Project-specific model weights update through lightweight fine-tuning while the foundation model remains frozen, eliminating the need to retrain billions of parameters. Data flows directly from SPR instruments to model training pipelines through automated processing, removing manual curation bottlenecks. Most critically, proprietary experimental data remains completely segregated—each project maintains isolated model weights that never intermingle with other projects’ data, ensuring pharmaceutical partners’ IP remains secure while still benefiting from the platform’s learning capabilities. The compound effect of rapid iteration becomes clear when comparing discovery timelines. In our chemokine receptor campaign, the ability to incorporate Round 1’s failures into Round 2’s design within 48 hours meant we could explore informed hypotheses while conventional approaches would still be synthesizing their initial round. This compression of learning cycles—from months to days—enables exploration of more diverse sequence space within the same project timeline, increasing the probability of finding optimal therapeutic candidates rather than settling for first acceptable hits. This rapid iteration capability aligns with the broader industry shift toward data-driven discovery paradigms [39, 119–124], but goes further by enabling true closed-loop optimization. Our case studies demonstrated a 3-4 fold reduction in experimental cycles—achieving in 3 rounds what traditional display methods require 8-12 rounds to accomplish. By progressing directly from computational design to recombinant validation without constructing extensive libraries, we eliminate months of library preparation while maintaining the ability to explore vast sequence diversity computationally. This represents not just acceleration but fundamental transformation in how therapeutic antibodies can be discovered and optimized.

### 6.5 Implementation Variants: Dual-Track Accessibility

#### Open-Source Implementation for Academic Research

The open-source implementation demonstrates our commitment to advancing the broader scientific community by providing a fully functional version for academic research. Built on ESM3 (esm3-open-2024-03), a 1.4B parameter variant of the state-of-the-art protein language model [99], this version delivers the core multi-shot learning capabilities that enabled our chemokine receptor success.

The implementation employs a sophisticated contrastive learning framework built on PyTorch, incorporating:

– **ESM3-based sequence encoding** with attention pooling mechanisms that identify relevant sequence regions
– **Contrastive projection head** with residual connections for improved gradient flow and training stability [125]
– **InfoNCE contrastive-loss function** for learning relative binding affinities between antibody-antigen pairs [126]
– **Parameter-efficient fine-tuning (PEFT)** using LoRA (rank = 16, α = 32) to adapt ESM3 with minimal memory footprint while maintaining performance

This academic version provides complete transparency for methodological validation and reproducibility. Researchers can examine the multi-shot learning architecture, validate our synthetic amplification approach, and contribute improvements through collaborative development. By providing comprehensive training materials and example datasets, we enable laboratories without extensive ML expertise to implement advanced computational antibody design. This democratization of access has already catalyzed innovations from multiple academic groups who have adapted the framework for their specific research challenges.

#### Commercial Platform for Therapeutic Development

The commercial platform (via abtique.com) extends beyond the open-source foundation with proprietary enhancements specifically designed for therapeutic development at scale. Advanced encoders incorporate hybrid sequence-structure representations when available, attention mechanisms tuned for long-range residue dependencies, and specialized modules for diverse antibody formats including bispecifics and nanobodies. These enhancements, developed through extensive pharmaceutical partnerships, provide the additional accuracy margins critical for clinical development decisions.

Delivered at abtique.com as a no-code platform, the commercial version eliminates technical barriers for antibody scientists. Users simply specify their target, antibody format, and any known constraints—the platform handles the entire computational workflow from sequence generation through multi-shot optimization. Automated integration with laboratory information management systems enables seamless data flow between lab assay results to model updates. Most critically for pharmaceutical partners, the platform guarantees complete data segregation, model training segregation, and potential IP exclusivity on generated sequences, ensuring competitive advantages remain protected.

#### Synergistic Impact of Dual Implementation

This dual-track approach creates synergistic benefits for the entire field. Academic researchers gain access to cutting-edge methods without licensing barriers, enabling them to push methodological boundaries and publish innovations that advance the science. Their findings and improvements flow back to enhance both versions. Meanwhile, pharmaceutical partners receive production-ready solutions with regulatory-compliant documentation, enterprise support, and IP protection necessary for therapeutic development.

The strategy reflects our belief that scientific progress requires both open collaboration and commercial innovation. By providing a robust academic version, we ensure our methods receive rigorous peer review and validation. By maintaining a premium commercial platform, we sustain the development resources necessary to continuously improve both versions. This balanced approach—rare in computational biology—accelerates progress across the entire antibody discovery ecosystem.

## 7. Conclusion

Hyperbind2 establishes multi-shot learning as a new paradigm for antibody discovery. Unlike static predictors that maintain constant success rates, our platform demonstrates progressive improvement through lab-to-AI feedback cycles, achieving 6.7× improvement within 3 rounds. This learning capability-enabled by the direct learning head model architecture and 24-48 hour update cycles fundamentally changes the economics and timelines of antibody discovery.

The convergence within 2-5 rounds, validated across multiple campaigns, provides a predictable path from initial screening to optimized candidates, which contrasts sharply with zero-shot approaches. The platform’s proven success -identifying 20 candidates ≤100 nM including 3 sub-10 nM binders -demonstrates that multi-shot learning can transform challenging targets from intractable to accessible.

Key contributions include the direct learning architecture that separates foundation representations from user-specific learning, the synthetic amplification methodology that generates O(n²) training signals from limited experimental data, and the establishment of short learning cycles that enable rapid iterative improvement. The platform’s ability to learn from conventionally discarded negative data provides crucial decision boundary information that static or zero-shot models fundamentally lack.

The dual implementation of HyperBind2 ensures broad impact: open-source democratizes access for academic research while the commercial platform provides accessible and accelerated capabilities for biologics research. By transforming antibody discovery from static prediction to progressive learning, Hyperbind2 addresses the fundamental limitation of zero-shot methods: their inability to adapt to novel, poorly characterized targets where therapeutic breakthroughs are most needed. This paradigm, where models improve through use rather than requiring massive retraining -represents a fundamental shift in how computational biology integrates with experimental workflows.

## 8. Software Availability and Data Accessibility

### Open-Source Implementation

Hyperbind2 is released under GPL-3.0 license. The GitHub repository contains full implementation based on ESM3 encoder, pre-trained weights, sample synthetic datasets, and comprehensive step-by-step training/inference scripts. All dependencies are listed in requirements.txt, and detailed hardware guidance is provided (≥ 16 GB GPU VRAM recommended; smaller cards supported with reduced batch size).

Repository: https://github.com/baverso/HyperBind2-OpenSource

The open-source release includes:

– Complete training and inference codebase with detailed documentation
– Pre-trained model weights for immediate use
– Synthetic dataset generation scripts and example datasets
– Comprehensive tutorials and example workflows
– Community support forums and contribution guidelines

### Commercial Platform

For access to the proprietary commercial version with enhanced capabilities and industrial-scale features, please visit abtique.com

## Data Availability

Synthetic datasets used in this study are available through the open-source repository. Proprietary experimental data from case studies are available upon reasonable request and subject to confidentiality agreements with collaborating partners.

## Reproducibility

All computational results presented in this paper can be reproduced using the provided open-source implementation and publicly available datasets. Detailed protocols and parameter settings are included in the repository documentation.

## Acknowledgements

We thank our experimental collaborators for providing the antibody screening data and validation measurements that comprise the case study presented here. Their rigorous experimental protocols and willingness to share both successful and unsuccessful selection rounds were essential for developing and validating our computational approach.

